# *Caenorhabditis elegans* PIEZO Channel Coordinates Multiple Reproductive Tissues to Govern Ovulation

**DOI:** 10.1101/847392

**Authors:** Xiaofei Bai, Jeff Bouffard, Avery Lord, Katherine Brugman, Paul W. Sternberg, Erin J. Cram, Andy Golden

## Abstract

The PIEZO proteins are involved in a wide range of developmental and physiological processes. Human PIEZO1 and PIEZO2 are newly identified excitatory mechano-sensitive proteins; they are non-selective ion channels that exhibit a preference for calcium in response to mechanical stimuli. To further understand the function of these proteins, we investigated the roles of *pezo-1*, the sole *PIEZO* ortholog in *C. elegans*. *pezo-1* is expressed throughout development in *C. elegans*, with strong expression in reproductive tissues. A number of deletion alleles as well as a putative gain-of-function mutant caused severe defects in reproduction. A reduced brood size was observed in the strains depleted of PEZO-1. *In vivo* observations show that oocytes undergo a variety of transit defects as they enter and exit the spermatheca during ovulation. Post ovulation oocytes were frequently damaged during spermathecal contraction. Calcium signaling in the spermatheca is normal during ovulation in *pezo-1* mutants, however, *pezo-1* interacts genetically with known regulators of calcium signaling. Lastly, loss of PEZO-1 caused defective sperm navigation after being pushed out of the spermatheca during ovulation. Mating with males rescued these reproductive deficiencies in our *pezo-1* mutants. These findings suggest that PEZO-1 may act in different reproductive tissues to promote proper ovulation and fertilization in *C. elegans*.

## Introduction

Mechanotransduction - the sensation and conversion of mechanical stimuli into biological signals - is essential for development. PIEZO1 and PIEZO2 are newly identified excitatory mechanosensitive proteins, which play important roles in a wide range of developmental and physiological processes in mammals (Alper, 2017; Coste et al., 2010; Coste et al., 2012; Murthy et al., 2017; Wu et al., 2017). PIEZO1 is a non-selective ion channel that forms homotrimeric complexes at the plasma membrane, however PIEZO1 exhibits a preference for Ca^2+^ in response to mechanical stimuli (Coste et al., 2010; Gnanasambandam et al., 2015; Syeda et al., 2015). Recent studies have shown that the human and mouse PIEZO1 channels respond to different mechanical stimuli, including static pressure, shear stress and membrane stretch (Coste et al., 2010; Poole et al., 2014; Ranade et al., 2014). PIEZO1 also regulates vascular branching and endothelial cell alignment upon sensing frictional force (shear stress) (Li et al., 2015; Nonomura et al., 2018). Stem cells also use PIEZO1 to sense mechanical signals and initiate Ca^2+^ signaling to promote proliferation and differentiation (Del Marmol et al., 2018; He et al., 2018). PIEZO2 primarily functions as a key mechanotransducer for light touch, proprioception and breathing (Nonomura et al., 2017; Woo et al., 2015; Woo et al., 2014). Mutations in both human *PIEZO1* and *PIEZO2* have been identified among the patients suffering from channelopathy diseases, such as Dehydrated Hereditary Stomatocytosis (DHS), Generalized Lymphatic Dysplasia (GLD), and Distal Arthrogryposis type 5 (DA5), in which osmoregulation is disturbed (Albuisson et al., 2013; Andolfo et al., 2013; Bae et al., 2013; Coste et al., 2013; Li et al., 2018; Lukacs et al., 2015; McMillin et al., 2014; Zarychanski et al., 2012). Loss-of-function mutations in the *PIEZO1* gene cause autosomal recessive congenital lymphatic dysplasia while gain-of-function mutations led to autosomal dominant stomatocytosis (Alper, 2017). However, the cellular and molecular mechanisms of PIEZO dysfunction in these diseases are not well understood.

*Caenorhabditis elegans* is an attractive model system to study mechanotransduction *in vivo. C. elegans* contains multiple tubular tissues, including the reproductive system, which experience mechanical stimulation throughout the life cycle (Cram, 2014, 2015; Voglis and Tavernarakis, 2005). The *C. elegans* reproductive system consists of two U-shaped gonad arms, each ending with a spermatheca and joined in the center by a shared uterus. *C. elegans* hermaphrodites produce sperm during the L4 larval stage and then shift to produce oocytes during the adult stage. About 150 sperm are stored in each spermatheca while the oocytes form in the oviduct in each gonad arm. The oocyte adjacent to the spermatheca undergoes oocyte maturation ∼25 minutes before being ovulated into the spermatheca (Greenstein, 2005). Oocyte maturation is triggered by sperm derived polypeptides known as major sperm proteins (MSPs), which activate the oocyte mitogen-activated protein kinase (MPK-1) (Miller, 2001; Yang et al., 2010). Once the oocyte matures, five pairs of contractile myoepithelial cells, named sheath cells, which make up the somatic gonad and encase the germline, push the matured oocyte into the spermatheca for fertilization. The spermatheca is an accordion-like multicellular tube, which consists of 28 myoepithelial cells surrounded by a basement membrane. Among the 28 cells, eight form the neck or distal valve of the spermatheca, four form the spermatheca-uterine valve that fuse into a syncytium, and 16 comprise a bag-like chamber between the two valves (Kimble and Hirsh, 1979; McCarter et al., 1999). The two spermathecal valves, distal (flanking the oviduct) and spermathecal-uterine (sp-ut) valve, are spatiotemporally coordinated to allow oocyte entry during ovulation and exit after fertilization, through acto-myosin contractions (Kelley and Cram, 2019). Ovulation is triggered by signaling between oocytes, sheath cells, and sperm through increasing cytosolic inositol 1,4,5-trisphosphate (IP3) and Ca^2+^ concentrations (Bui and Sternberg, 2002; Clandinin et al., 1998; Han et al., 2010). The ovulated oocyte spends 3-5 minutes in the dilated spermatheca with both valves closed to allow the oocyte and sperm to complete fertilization and to initiate eggshell formation. The constriction of the spermathecal bag cells and the opening of the spermathecal-uterine valve cells expel the fertilized egg into the uterus. Meanwhile, the sperm that are swept out of the spermatheca during oocyte exit crawl back to the constricted spermatheca. The navigation of the sperm back to the spermatheca is regulated by the chemoattractant prostaglandin that is secreted by the oocytes and sheath cells (Kubagawa et al., 2006). The signaling during spermathecal transit requires increased levels of IP3 and cytosolic Ca^2+^, which is regulated by filamin FLN-1, phospholipase PLC-1, and the endoplasmic reticulum calcium release channel ITR-1 (Kovacevic et al., 2013). Despite the probable role of mechanical stimuli during this whole process, such as stretch of oocyte entry or the contraction of the spermatheca, the mechanisms underlying the mechanosensitive channels in ovulation and fertilization remains largely unknown.

Unlike most vertebrates that have two *PIEZO* genes, only one ortholog, *pezo-1*, has been identified in the *C. elegans* genome. *Drosophila* also has a single PIEZO (Coste et al., 2012; Kim et al., 2012). A total of 14 different transcriptional isoforms of the *C. elegans pezo-1* gene have been identified, which are predicted to encode 12 different PEZO-1 proteins ranging from 1038 to 2442 amino acids. All of the isoforms share a common C-terminal domain that is highly conserved with both human PIEZO1 and PIEZO2. In this study, we used CRISPR/Cas9 gene editing, genetic approaches, and high-resolution confocal microscopy to characterize the function of PEZO-1 in *C. elegans*. We demonstrate a global expression pattern of PEZO-1 in different tubular tissues, including the reproductive system. We hypothesized that a mechanosensitive protein such as PEZO-1 would be involved in processes that involve such cellular movements as those involved in ovulation, where oocytes must transit into and out of the spermatheca. Multiple deletion mutations, as well as a putative gain-of-function mutation, caused severe reproductive deficiencies, such as reduced brood size, and defects in ovulation and sperm navigation. To examine whether PEZO-1 influences cytosolic Ca^2+^ signaling, imaging using the calcium indicator GCaMP3 revealed normal calcium release in the spermatheca during ovulation of *pezo-1* mutants compared with wild type. However, depletion of Ca^2+^ channels *itr-1* and *orai-1*, and Ca^2+^ transporting ATPase *sca-1*, by RNAi substantially enhanced the reproductive deficiencies in *pezo-1* mutants. The *pezo-*1 mutants displayed a sperm navigational defect as well as a reduced ovulation rate. Sperm that were readily washed out of the spermatheca during ovulation failed to migrate back to the spermatheca, thus depleting the spermatheca of sperm early in the reproductive lifecycle. Supplementing male sperm via mating significantly repopulated the spermatheca with sperm, and rescued the extremely low ovulation rate and reduced brood size in *pezo-1* mutants.

Using an auxin-inducible degradation (AID) system, we depleted PEZO-1 in different tissues, including somatic tissues, sperm, and the germline. Reduced brood sizes were observed in each tissue-specific degradation strain, suggesting multiple inputs of PEZO-1 from many tissues in regulating reproduction. To our knowledge, this is the first report of a functional role for PEZO-1 in *C. elegans* reproduction.

## Results

### PEZO-1 is expressed in multiple tissues throughout development

The *C. elegans* genome encodes a single *PIEZO* ortholog, *pezo-1*, of which there are 14 mRNA isoforms due to differential splicing and transcriptional start sites (Fig. S1A) (Harris et al., 2019); these 14 isoforms code for 12 different PEZO-1 proteins. All isoforms share a common C-terminus. To accurately visualize the expression pattern of *pezo-1 in vivo*, we directly knocked-in different fluorescent reporter genes into both the N-terminus and C-terminus of the *pezo-1* endogenous locus using CRISPR/Cas9. The C-terminal knock-in reporters should tag all *pezo-1* isoforms, while the N-terminal knock-in reporters should only tag the eight longest *pezo-1* isoforms (Fig.1A, S1A). Both GFP and mScarlet were used as reporters to generate N- and C-terminal fusions proteins. GFP::PEZO-1, mScarlet::PEZO-1, and PEZO-1::mScarlet were widely expressed from embryonic stages through adulthood (Fig. 1B-I, Fig. S1B-G). The genome-edited animals behaved normally, suggesting no functional disruption of tagging PEZO-1 with these fluorescent reporter genes. Notably, PEZO-1 is strongly expressed in several tubular tissues, including the pharyngeal-intestinal and spermathecal-uterine valves, which is consistent with our hypothesis that *pezo-1* may be responsible for mechanoperception in these tissues (Fig. 1B, Fig. S1B, C). Under higher magnification, we observed PEZO-1 on the plasma membranes of oocytes and embryonic cells during a variety of embryonic stages, suggesting PEZO-1 is a transmembrane protein (Fig. 1C-H). Partial co-localization of PEZO-1 with the ER marker SP12::GFP suggested that PEZO-1 may be processed in the ER and transported to the plasma membrane (Fig. 1E). PEZO-1 is expressed in multiple reproductive tissues, including the germline, somatic oviduct, and spermatheca (Fig. 1F-H). Higher magnification imaging of the spermatheca revealed expression of PEZO-1 on sperm membranes as well (Fig. 1I). Consistent with the hypothesis that reproductive tissues are regulated by mechanosensitive stimuli in *C. elegans*, expression of PEZO-1 likely functions to sense physical strain or contractility during ovulation and fertilization. Live imaging and detailed analysis of PEZO-1 expression patterns during reproduction revealed that GFP::PEZO-1 is expressed in sheath cells, sperm, both spermathecal valves and the spermathecal bag cells (Fig. 1J-N, Video S1). The fluorescent signal of GFP::PEZO-1 is observed in both spermathecal valves until the valves open, suggesting that PEZO-1 may function to sense the mechanical stimuli at the valves during ovulation (Fig. 1J, L, M, Video S1). As the fertilized oocyte is pushed into the uterus, GFP::PEZO-1 labeled sperm crawl back into the constricting spermatheca after each ovulation (Fig. 1N, Video S1). Collectively, these data indicate that PEZO-1 is expressed in the somatic gonadal cells and germline cells. To our knowledge this is the first observation of a mechanosensitive channel in the process of ovulation in *C. elegans*.

**Figure 1.**
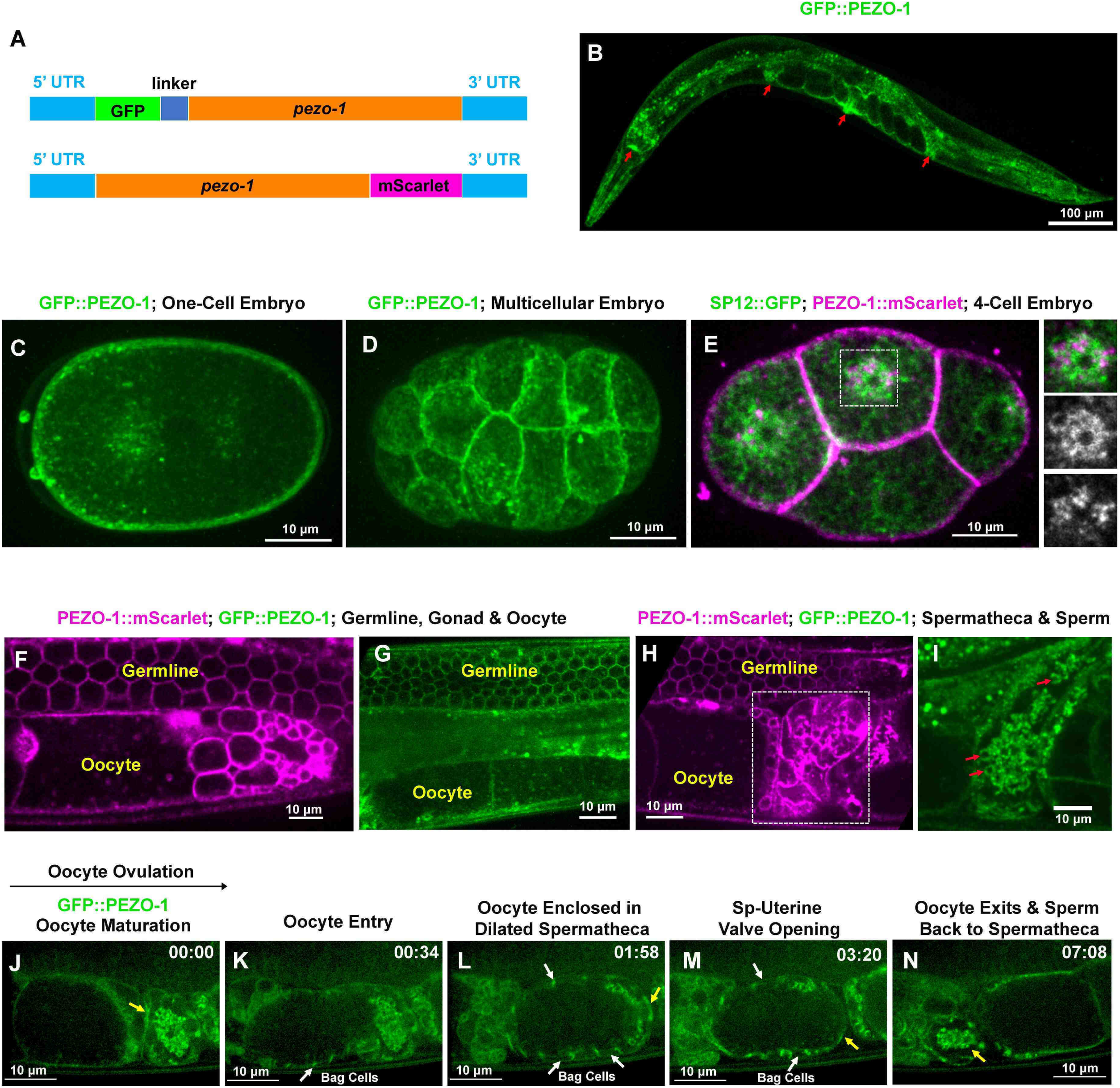
*pezo*-1 expression pattern in *C. elegans*. *pezo-1* is globally expressed in *C. elegans*. (A) Two fluorescent reporter genes were knocked-in to both the *pezo-1* N-terminus and C-terminus. (B) GFP::PEZO-1 is strongly expressed in multiple mechanosensitive tissues, such as the pharyngeal-intestinal valve, spermatheca, and vulva (red arrows). (C, D) GFP::PEZO-1 (green) is expressed in the plasma membrane of different staged embryos. (E) PEZO-1::mScarlet (magenta) localizes to the plasma membrane and a small fraction co-localizes with the ER marker SP12::GFP (green). Right insets show PEZO-1::mScarlet (bottom panel), SP12::GFP (middle panel), and merged image (top panel) on the ER. (F-I) Both PEZO-1::mScarlet (magenta) and GFP::PEZO-1 (green) localize to reproductive tissues, such as the plasma membrane of the germline cells (F-H), somatic gonad (F-H), spermatheca (H; in white box), and sperm (red arrows) (I). (J-N) Representative images of PEZO-1 localization during ovulation and fertilization. GFP::PEZO-1 (green) localizes to the spermathecal distal valve (yellow arrow, J), which remains closed before ovulation. The oocyte ovulated and entered into the spermatheca (K) and stayed enclosed in the spermatheca until fertilization completed (L). During fertilization, GFP::PEZO-1 remained on the spermathecal-uterine (sp-ut) valve as indicated by a yellow arrow (L, M). The bag cells of the spermatheca also express GFP::PEZO-1 at this time (representative bag cells are marked by white arrows, K-M). After fertilization, the sp-ut valve opened (M, yellow arrow) and allowed the newly-fertilized zygote to exit the constricting spermatheca (M, N). Once a fertilized zygote entered the uterus, the spermatheca constricts; sperm can be seen in the constricted spermatheca (N, yellow arrow). Black arrow above panel J shows the direction the embryo travels through the spermatheca from left to right. Timing of each step is labeled on the top right in minutes and seconds. Scale bars are indicated in each panel.

### Deletion of *pezo-1* causes a decrease in brood size

To further investigate the function of *pezo-1*, the phenotypes of *pezo-1* knockout (*pezo-1*^KO^) animals were analyzed. Two candidate null alleles were generated by CRISPR/Cas9 genome editing; one allele was a deletion of exons 1-13 (*pezo-1 N-Δ*) and the other bearing a deletion of the last seven exons, 27-33 (*pezo-1 C-Δ*) (Fig. S2A-B). Two other alleles were generated by CRISPR/Cas9: *pezo-1 (sy1398)*, which has a deletion of an exon unique to the two shortest isoforms, i and j, and a putative null allele, *pezo-1 (sy1199)*, which has a “STOP-IN” mutation in exon 27 that should interfere with translation of the C-termini of all isoforms (Fig. S2B). Although GFP::PEZO-1 and PEZO-1::mScarlet expressed globally in adult worms, we did not observe obvious morphological phenotypes from homozygous *pezo-1^KO^* mutants compared to control animals. However, in all tested *pezo-1* mutants, the number of F1 progeny was significantly reduced compared with wild type (Fig. 2A, S2C). The decrease in brood size was enhanced as animals aged (36-60 hours post mid-L4, Fig. 2B, S2C) or when grown at a higher temperature (25°C, Fig. S2D). In addition, about 5-25% of F1 embryos failed to hatch from *pezo-1 C-Δ* homozygous mutants (Fig. 2B). To mimic a gain-of-function phenotype in *pezo-1*, we fed wildtype animals with Yoda1, a PIEZO1 specific chemical agonist, which keeps the channel open (Syeda et al., 2015). Reduced brood sizes were observed when wildtype animals were exposed to 100 and 200 μM Yoda1 (Fig. 2C). These data suggest that either deletion or overactivation of PEZO-1 is sufficient to disrupt brood size.

**Figure 2.**
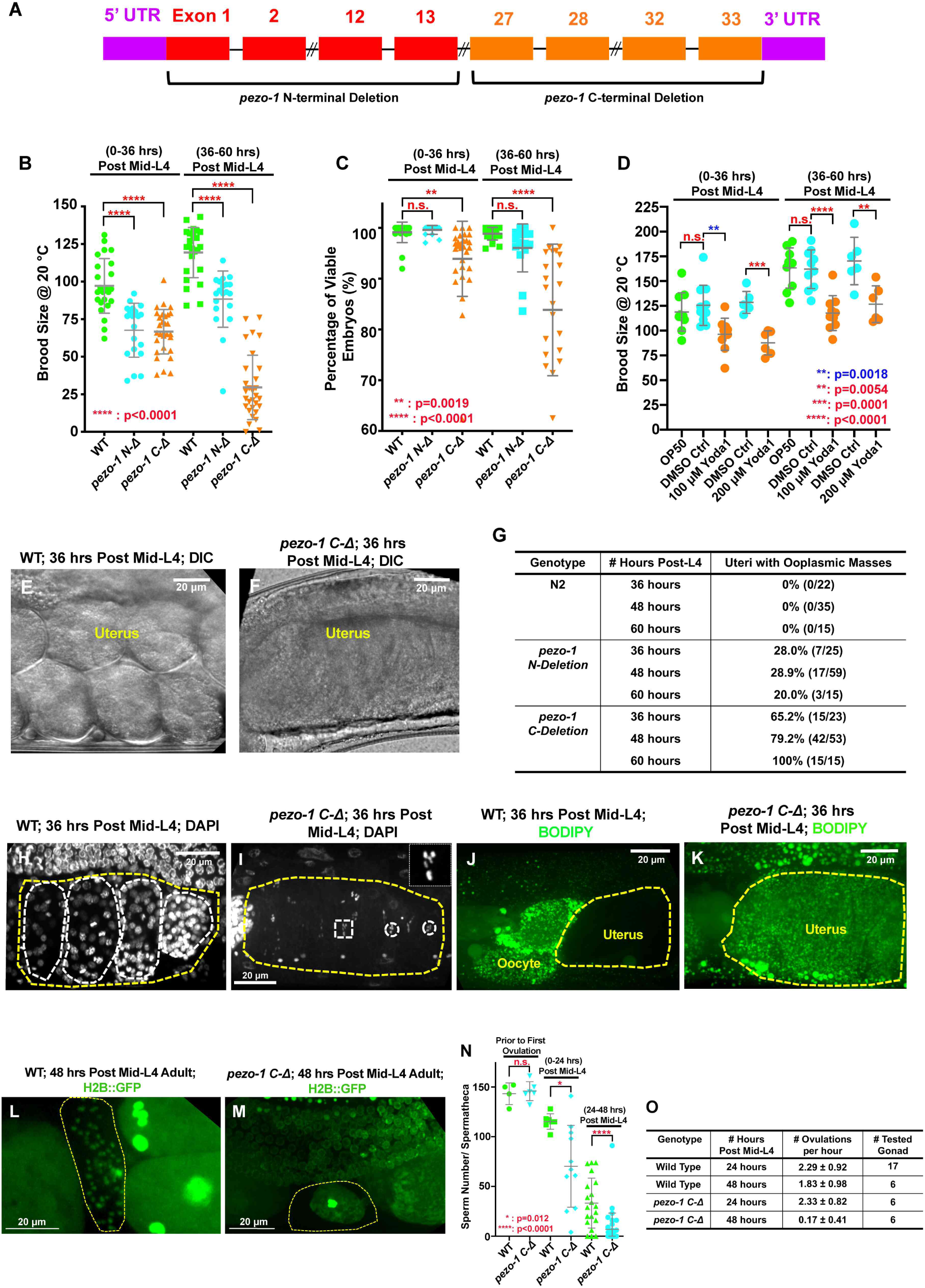
Deletions of the *pezo-1* gene cause a reduction in brood size. (A) Brood size was significantly reduced in both *pezo-1 N-Δ* and *pezo-1 C-Δ* animals when compared with wildtype and this reduction was most evident in older adult animals. (B) The percentage of viable embryos was reduced in the *pezo-1 C-Δ* animals. (C) Dietary supplementation of a PIEZO1 channel specific activator Yoda1 in wildtype animals significantly reduced the brood size compared with control treatment. (D, E) DIC images of the uteri of gravid adult animals. Wildtype animals had young embryos in their uteri (D), while only a large ooplasmic mass was observed in *pezo-1 C-Δ* mutant uteri (E). (F) Quantification of the percentage of uteri with ooplasmic masses in wildtype and *pezo-1* deletion mutants. N2 is the wildtype strain. (G, H) DAPI staining demonstrated that multicellular embryos (white circles, G) were present in the uteri of wildtype animals, while only oocyte meiotic chromosomes (white circles and rectangle) were observed in the uteri of *pezo-1 C-Δ* mutants (H; inset in top right white box shows an amplified image of the meiotic chromatin marked with a white rectangle). The yellow dotted lines indicate the boundaries of the uteri in panels G and H. (I, J) Only unfertilized oocytes and newly-fertilized zygotes are permeable to BODIPY (green) in wildtype (WT) animals (I), while staining was observed throughout the entire uterine mass (yellow circle, J) of *pezo-1 C-Δ* animals. (K, L) An H2B::GFP transgene was crossed into our strains to visualize oocyte and sperm chromatin. (K) Sperm labeled by H2B::GFP (green cells in yellow circle) reside in the spermatheca (yellow circle) of day 2 adults (48 hours post mid-L4). (L) Only oocyte debris (yellow circle) is left in the spermatheca of an age-matched *pezo-1 C-Δ* mutant. (M) Quantification of sperm counts in both wild type and *pezo-1 C-Δ* hermaphrodites at different time windows. (N) Quantification of the oocyte ovulation rate of wildtype and *pezo-1 C-Δ* adults at different ages. The oocyte ovulation rate was significantly reduced in the older *pezo-1 C-Δ* mutant adults. P-values: * = 0.012 (M); ** = 0.0019 (B); ** = 0.0018 (C, blue); ** = 0.0054 (C, red); *** =0.0001 (C); **** <0.0001 (*t*-test).

Next, using differential interference contrast (DIC) and confocal microscopy, we analyzed the defects associated with the observed reduction in brood size. While embryos fill the uterus in wildtype mothers (Fig. 2D), a mass of ooplasm in the uteri of both *pezo-1^KO^* and stop-in mutants was observed (Fig. 2E, S2E). Occasionally, a few fertilized embryos were observed inside this mass of ooplasm (data not shown). *pezo-1 C-Δ* and STOP-IN mutants displayed the most severe defects, where 100% of animals had a uterus filled with ooplasm at 60 hours post L4 (Fig. 2F, S2E), Staining with DAPI in *pezo-1^KO^* uteri revealed chromosome structures indicative of diakinesis-staged oocytes (Fig. 2H). In contrast, only mitotic chromatin of variably-aged embryos were detected in control animals (Fig. 2G). Consistent with this observation, only unfertilized oocytes and newly-fertilized embryos without intact eggshells stained with the lipophilic dye, BODIPY, in wildtype animals (Fig. 2I). BODIPY staining revealed widespread penetration of the entire ooplasmic mass in the uteri of *pezo-1 C-Δ* animals (Fig. 2J). These data suggest that oocytes are not fertilized upon transit through the spermatheca and that these unfertilized oocytes may be crushed when they pass through the spermathecal valves. A more detailed characterization of the ovulation defects is described in the next section.

In addition to these apparent crushed oocytes, a reduced number of sperm resident in the spermatheca were observed in day 1 *pezo-1* adults (0-24 hours post mid-L4) and even fewer were observed in the spermathecae in day 2-3 adults (24-48 hours post mid-L4) compared with wild type (Fig. K-M). Normal numbers of sperm were present in these mutant hermaphrodites prior to the first ovulation, suggesting that the ability of the sperm to return to the spermatheca after each ovulation was disrupted (Fig. 2M). Sperm loss could also contribute to the low brood sizes observed in our *pezo-*1 mutants.

Ovulation rates were significantly reduced in *pezo-1 C-Δ* day 2 (post mid-L4 48 hours) animals (Fig. 2N), which is consistent with the reduced brood sizes that worsen in day 2 animals. Since sperm presence in the spermatheca is known to stimulate ovulation (McCarter et al., 1999; Miller, 2001), the reduction in sperm number could be responsible for this reduction in ovulation rate. Overall, the reduced brood size in *pezo-1* mutants is likely due to the consequential defects in multiple tissues, resulting in defective ovulations, crushed oocytes, and defects in the ability of sperm to navigate back into the spermatheca after each ovulation.

### Severe ovulation defects were observed in the *pezo-1* mutants

To carefully characterize the transit of oocytes through the spermatheca, we performed live imaging to record the ovulation and fertilization process in both wildtype and *pezo-1^KO^* animals (Fig. 3 A-E’, Video S2-3). The imaging began with the mature oocyte entering the spermatheca, labeled by the apical junction marker DLG-1::GFP (Fig. 3A, B). In wildtype animals, the contracting sheath cells push the oocyte into the spermatheca, and simultaneously pull the open spermatheca over the oocyte (Video S2-3). Once the oocyte enters the spermatheca, both spermatheca valves remain closed during fertilization (Fig. 3C). Opening of the sp-ut valve allows the fertilized oocyte to be expelled into the uterus (Fig. 3D, E). However, in *pezo-1* mutants, we frequently observed that oocytes partially entered the spermatheca but were then pinched off and broken into two pieces, one of which remained trapped in the oviduct (proximal gonad; Fig. 3F-I, Video S4). Moreover, some oocytes failed to enter the spermatheca and slid back into the oviduct (Fig. 3J-M, Video S5). The defective ovulation is likely due to incomplete constriction of the sheath cells and improper gating of the distal spermathecal valve. Of the oocytes that did successfully enter the spermatheca, many were frequently crushed when they exited through the sp-ut valve (Fig. 3A’-E’, Video S2-3). We observed that the sp-ut valve, labeled by DLG-1::GFP, did not completely open when the oocyte attempted to exit the spermatheca, which may lead to crushing the oocyte (Fig. 3C’-E’, Video S3). The ooplasm from the crushed oocytes accumulated in the uterus (Fig. 3E’, Video S3) as a large ooplasmic mass (as shown in Fig. 2E). Overall, disrupted ovulation and oocyte transit defects were observed in *pezo-1* mutants, consistent with the decreased brood size observed in all of our *pezo-1* mutants.

**Figure 3.**
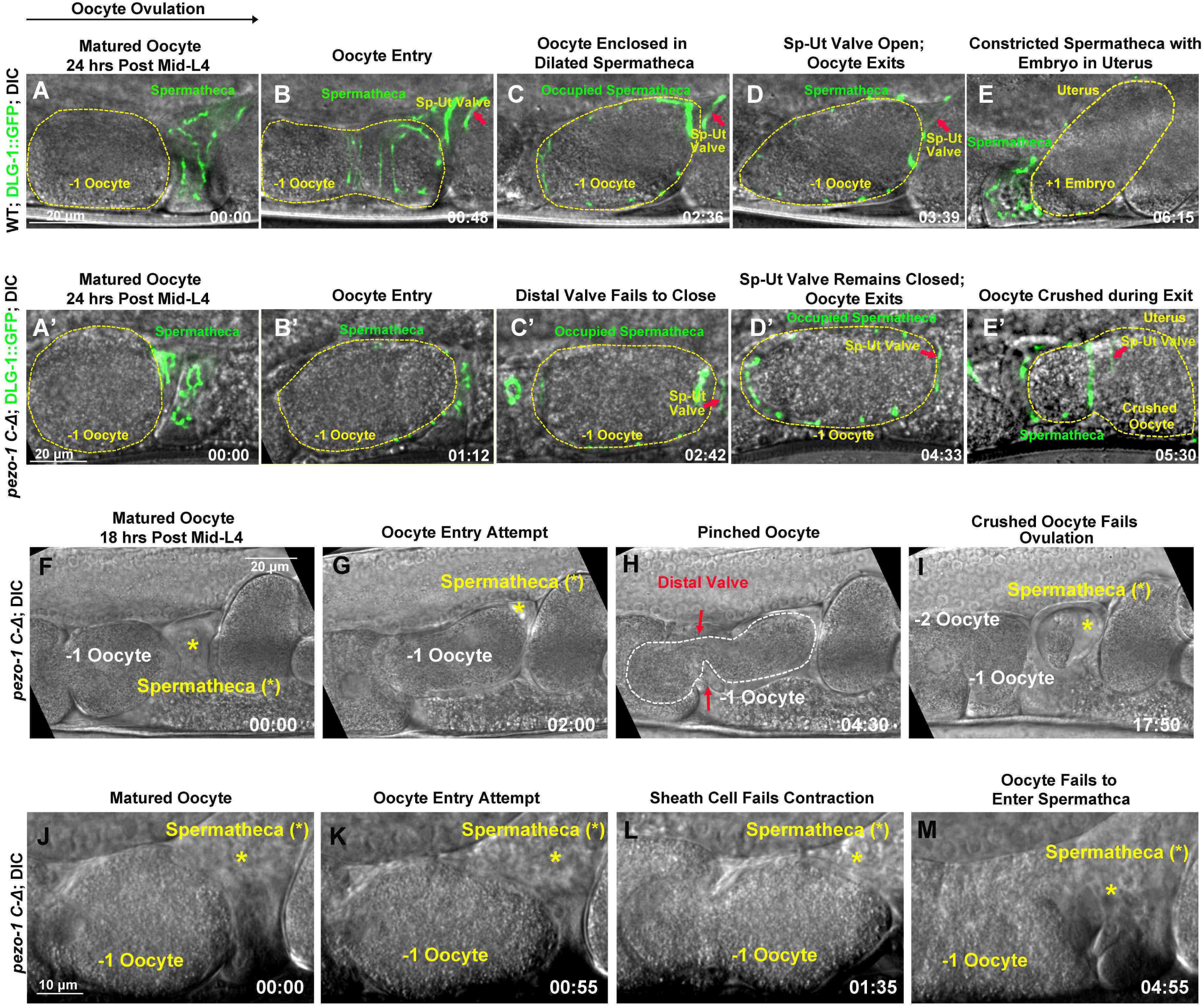
PEZO-1 mutants cause severe ovulation defects. (A-E) Ovulation in wildtype animals. (A, B) Ovulation is initiated by oocyte (yellow dotted circle) entry into the spermatheca, which was labelled by the apical junctional marker DLG-1::GFP (green). (C) Fertilization occurs in the occupied spermatheca (yellow dotted circle). (D-E) After fertilization, the sp-ut valve (red arrows) opened immediately to allow the newly-fertilized zygote (yellow dotted circle) to exit the spermatheca and enter the uterus. (A’-E’) Abnormal ovulation was observed in *pezo-1 C-Δ* animals. Control of the spermathecal valves was aberrant (C’-E’) during ovulation and the DLG-1::GFP labelled sp-ut valve (red arrow) never fully opened; the oocyte was crushed as it was expelled (E’). (F-M) Two examples of failed ovulations. (F-I) Ovulation defects observed in the *pezo-1 C-Δ* mutants. The ovulating oocyte (white dotted circle) was pinched off by the spermathecal distal valve (red arrows, H). This oocyte never exited into the uterus. (J-M) *pezo-1 C-Δ* oocyte frequently failed to enter the spermatheca and was retained in the oviduct (M). Black arrow above panel A shows the direction the embryo travels through the spermatheca from left to right. Timing of each step is labeled on the bottom right in minutes and seconds. Scale bars are indicated in each panel.

### PEZO-1 mutants genetically interact with cytosolic Ca^2+^ regulators

Given that PEZO-1 is the ortholog of mammalian mechanosensitive calcium channels and may regulate spermathecal contractility, in *C. elegans*, through Ca^2+^ signaling pathways, we tested whether there were genetic interactions between *pezo-1* mutants and several essential cytosolic Ca^2+^ regulators. Firstly, ITR-1, an ER Ca^2+^ release channel and an inositol-1,4,5-triphosphate (IP3) kinase LFE-2 were depleted by RNAi in both wildtype and *pezo-1* mutants. IP3 binding to ITR-1 releases Ca^2+^ from the ER, which activates myosin for spermathecal contractility (Bouffard et al., 2019; Clandinin et al., 1998; Kovacevic et al., 2013). Therefore, we hypothesized that combining *pezo-1* mutants with *itr-1* RNAi would greatly exacerbate the reduction in brood size if they were both critical to ovulation and fertilization. We carefully calibrated *itr-1* RNAi treatment and determined that feeding L4 animals for 36-60 hours produced optimal intermediate conditions that caused minimal developmental defects and normal brood sizes in wildtype animals. Consistent with our hypothesis, feeding *itr-1* RNAi enhanced the reduction of brood size of *pezo-1* mutants compared with wild type (Fig. 4A). In contrast, feeding *lfe-2* RNAi, which should elevate cytosolic Ca^2+^, partially rescued the reduced brood size (Fig. 4B). Therefore, *pezo-1^KO^* mutants have a negative genetic interaction with *itr-1*(RNAi) and a positive genetic interaction with *lfe-2* (RNAi). Similarly, depletion of the plasma membrane Ca^2+^ channel *orai-1*, which is activated to replenish the extracellular Ca^2+^ to the cytosol (Lorin-Nebel et al., 2007), led to nearly zero brood size in *pezo-1 C-Δ* mutant but only a 40% reduction in brood size in wild type (Fig. 4C). Furthermore, disruption of ER Ca^2+^ stores with sarcoplasmic/ER Ca^2+^ ATPase (SERCA) *sca-1*(RNAi) (Yan et al., 2006) also caused an extremely low brood size in *pezo-1 C-Δ* (Fig. 4C) while *sca-1*(RNAi) slightly increased the brood size in wild type (Fig. 4C). Therefore, these observations are consistent with the hypothesis that *pezo-1* may regulate cytosolic and ER Ca^2+^ homeostasis, which is crucial for proper spermathecal contractility and dilation.

**Figure 4.**
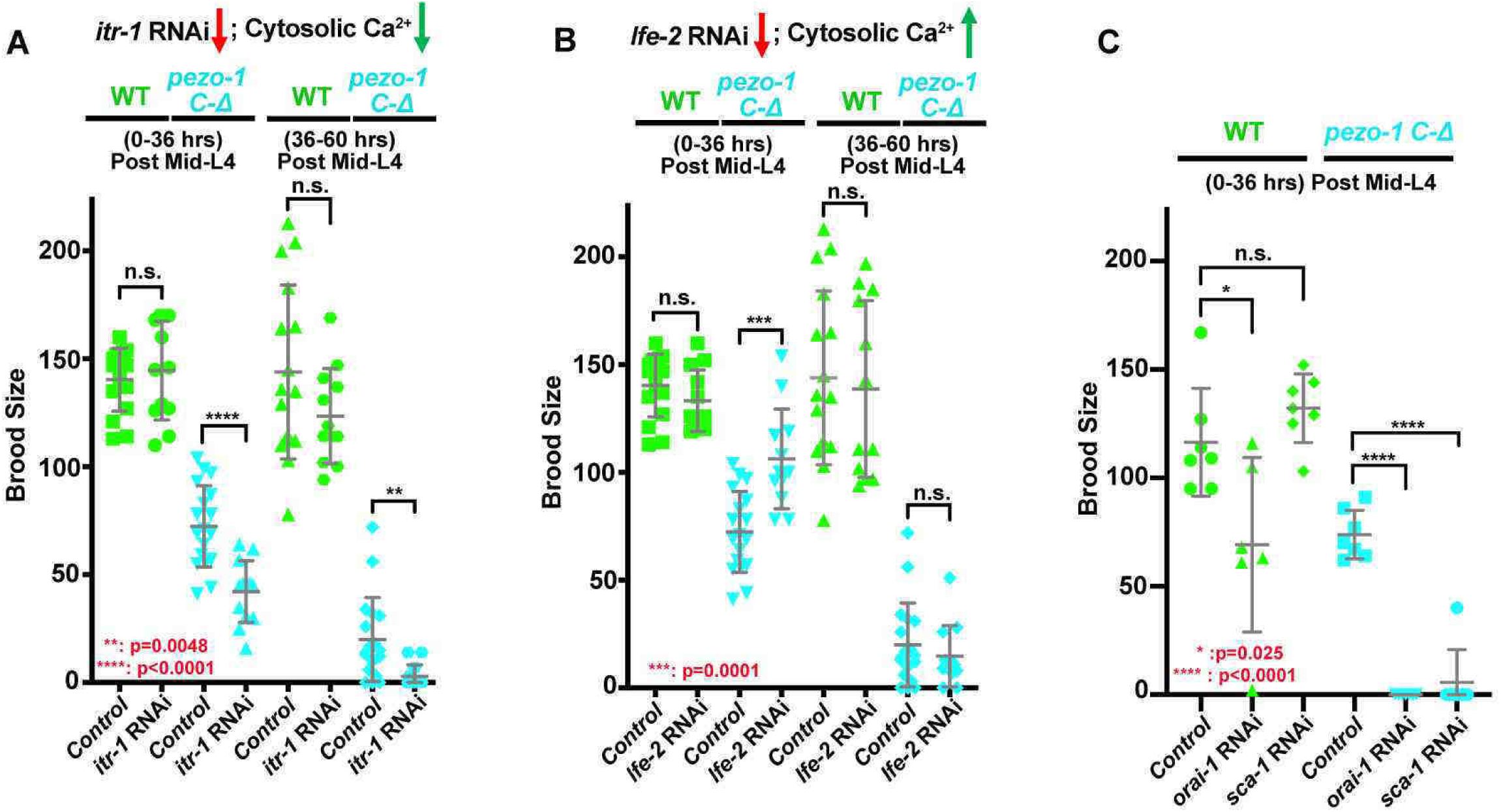
*pezo-1* mutants genetically interact with cytosolic Ca^2+^ regulators. (A) *itr-1 (RNAi)* reduced the brood size in *pezo-1 C-Δ* animals. (B) In contrast, *lfe-2 (RNAi)* slightly rescued the smaller brood size in *pezo-1 C-Δ* animals. (C) Depletion of both *orai-1* and *sca-1* by RNAi also enhanced the brood size reduction of *pezo-1 C-Δ* mutants. P-values: * = 0.025 (C); ** = 0.0048 (A); *** = 0.0001 (B); **** <0.0001 (*t*-test).

### *pezo-1* mutants show normal calcium signaling in spermatheca cells during ovulation

Due to PIEZO channels’ permeability to Ca^2+^ and the importance of calcium signaling in regulating spermathecal contractility, we tested whether deletion of *pezo-1* disrupted cytosolic Ca^2+^ homeostasis. We imaged oocyte passage through the spermatheca of both wild type and *pezo-1* mutants expressing the Ca^2+^ indicator GCaMP3, which was driven by a spermathecal-specific *fln-1* promoter (Bouffard et al., 2019; Kovacevic et al., 2013). Co-localization of the GCaMP3 transgene with mScarlet::PEZO-1 in the spermatheca suggested that this transgene would be useful for the analysis of *pezo-1* function in spermathecal calcium signaling (Fig 5A-E, Video S6). To determine whether calcium signaling was altered in our *pezo-1* mutants, a set of high-speed GCaMP imaging data from different animals was generated and the average pixel intensity of each frame was quantified (Fig 5F-J’, S3A-D, Video S6). We defined the initial time frame as the time just before the oocyte entered the spermatheca. In wildtype animals, the fluorescent intensity of GCaMP3 at the distal valve immediately increased when the oocyte entered the spermatheca (Fig 5A, F and F’, Video S6-7). During fertilization, an increase in intensity of GCaMP3 was frequently observed in the bag cells and sp-ut valve until the oocyte exited the spermatheca (Fig 5B-D, G-I and G’-I’, Video S6-7). GCaMP3 signal decreased to basal intensity after the fertilized oocyte was expelled into the uterus (Fig. 5E, J and J’, Video S6-7). To statistically quantify and analyze the oocyte transit, we defined a series of parameters, including the dwell time and two calcium signaling metrics from the GCaMP3 time series (Bouffard et al., 2019). A spermathecal tissue function metric, dwell time, is defined as the time from spermathecal distal valve closure to sp-ut valve opening, which represents the duration of time that the oocyte resides in the enclosed spermatheca. The calcium signaling metric, fraction over half max, is defined as the duration of the dwell time over the GCaMP3 half-maximal value divided by the total dwell time. The fraction over half max allows us to capture the relative level of calcium throughout the time the embryo passes through the spermatheca. Rising time indicates the time from the opening of the distal valve to the first time point at which the GCaMP fluorescent intensity reaches half maximum (Bouffard et al., 2019). In *pezo-1 C-Δ* mutants, longer transit times of the oocyte through the spermatheca resulted in elongated dwell times (Fig. 5K, Video S7), suggesting that deletion of *pezo-1* resulted in disrupted tissue function. Surprisingly, GCaMP3 fluorescence in *pezo-1* was not significantly different than wildtype (Fig. 5L, M, Video S7; see methods). GCaMP3 time series (Fig. S3A, B, Video S7), heat maps (Fig. S3C), and kymograms (Fig. S3D, E) also displayed normal Ca^2+^ levels during oocyte passage through the spermatheca in *pezo-1* mutants. It should be noted that we only imaged the GCaMP3 reporter during the very first three ovulations in young adult animals to avoid Ca^2+^ signaling interference from a distorted gonad morphology and mechanical pressure from a gravid uterus. Furthermore, it is difficult to monitor older hermaphrodites as they do not ovulate well on microscope slides. Since only mild defects were observed in the *pezo-1* mutants during these early ovulations and oocyte transit defects increased in severity over time (Fig. 2F), our data does not exclude the possibility that Ca^2+^ signaling may be more severely disrupted as the animal goes through more ovulation cycles. Alternatively, the live imaging assay may not be sensitive enough to detect subtle variations in calcium signaling.

**Figure 5.**
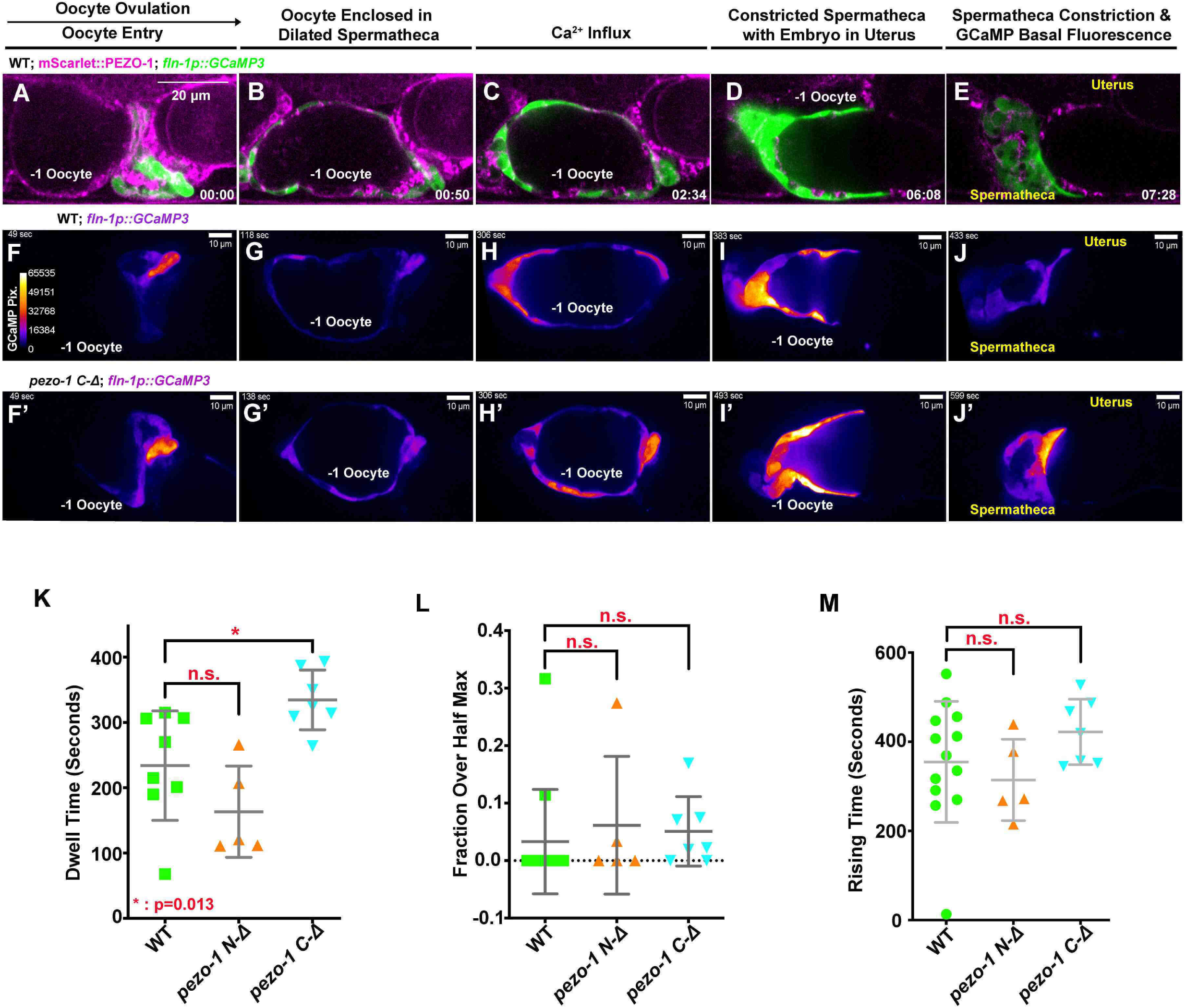
PEZO-1 mutants show normal GCaMP3 fluorescence during ovulation. (A-E) mScarlet::PEZO-1 colocalizes with GCaMP3 that is driven by a spermatheca-specific promoter. These images represent the third ovulation for this animal. (F-J’) Time series frames from GCaMP3 recordings in the third ovulation of both wildtype animals (F-J) and *pezo-1 C-Δ* animals (F’-J’). Ca^2+^ influx was quantified during ovulation and fertilization, as indicated by the intensity of GCaMP3 pixels (colored bar in F). (F, F’) Oocyte entry into the spermatheca in wild type and *pezo-1 C-Δ.* (G, G’) Oocytes in the spermatheca, (H, H’) Ca^2+^ influx during fertilization, (I, I’) intense Ca^2+^ influx as sp-ut valve closes to push newly-fertilized zygote into the uterus, and (J, J’) the return to basal levels as the spermatheca prepares for the next ovulation. (K) Dwell time is a tissue function metric calculated as the time the oocyte resides in the spermatheca from the closing of the distal valve to the opening of the sp-ut valve. (L, M) Calcium signaling metrics, fraction over half max (L), rising time (M) in *pezo-1* mutants showed normal calcium levels during ovulation compared with wild type. Black arrow above panel A shows the direction the embryo travels through the spermatheca from left to right. Timing of each step is labeled on the bottom right in minutes and seconds (A-E). Scale bars are indicated in each panel.

### Sperm from matings rescues low brood size phenotype in *pezo-1* mutants

In *C. elegans*, successful ovulation and fertilization requires signal coordination between sperm, oocytes, and sheath cells (Han et al., 2010). Given that PEZO-1 is expressed in these tissues, it is plausible that oocyte transit defects and reduced brood sizes are due to impaired inter-tissue signaling, which may be mediated by PEZO-1. To investigate how this may occur, bidirectional signaling between sperm and oocytes was first tested. To test sperm fertility, both wildtype and *pezo-1* mutant males were mated with *fem-1(hc17)* hermaphrodites, which do not produce any sperm or self-progeny (Doniach and Hodgkin, 1984) and are essentially females. The *fem-1(hc17)* animals produced cross-progeny after mating with *pezo-1* mutant males, indicating *pezo-1* mutant males are fertile and that their sperm can crawl through the uterus to the spermatheca upon mating (Fig. 6A). Since *pezo-1* mutant hermaphrodites do not produce any self-progeny after day 3 (60 hours post mid-L4) (Fig. 6B), we tested whether mating with both wildtype and mutant males would result in any cross progeny in the aged *pezo-1* mutants. *pezo-1* mutant hermaphrodites resumed ovulation and fertilization upon mating once the male’s sperm (from either wildtype or *pezo-1* males) reached the spermatheca (Fig. 6B-D). To test whether sperm signaling was defective in *pezo-1* mutants, we mated *spe-9(hc52ts)* males with both wildtype and *pezo-1* mutant hermaphrodites. *spe-9(hc52ts)* male sperm can physically contact the oocytes but fails to fertilize, however, the sperm signaling is apparently normal and triggers ovulation (Singson et al., 1998). Interestingly, the low ovulation rate in *pezo-1 C-Δ* animals was significantly rescued by *spe-9(hc52ts)* sperm (Fig. 6E), although the ovulated oocytes were not fertilized. Overall, these data suggest that the absence of sperm contributes to a profound reduction of oocyte maturation, ovulation rate, and self-fertility in the aged *pezo-1* mutants.

**Figure 6.**
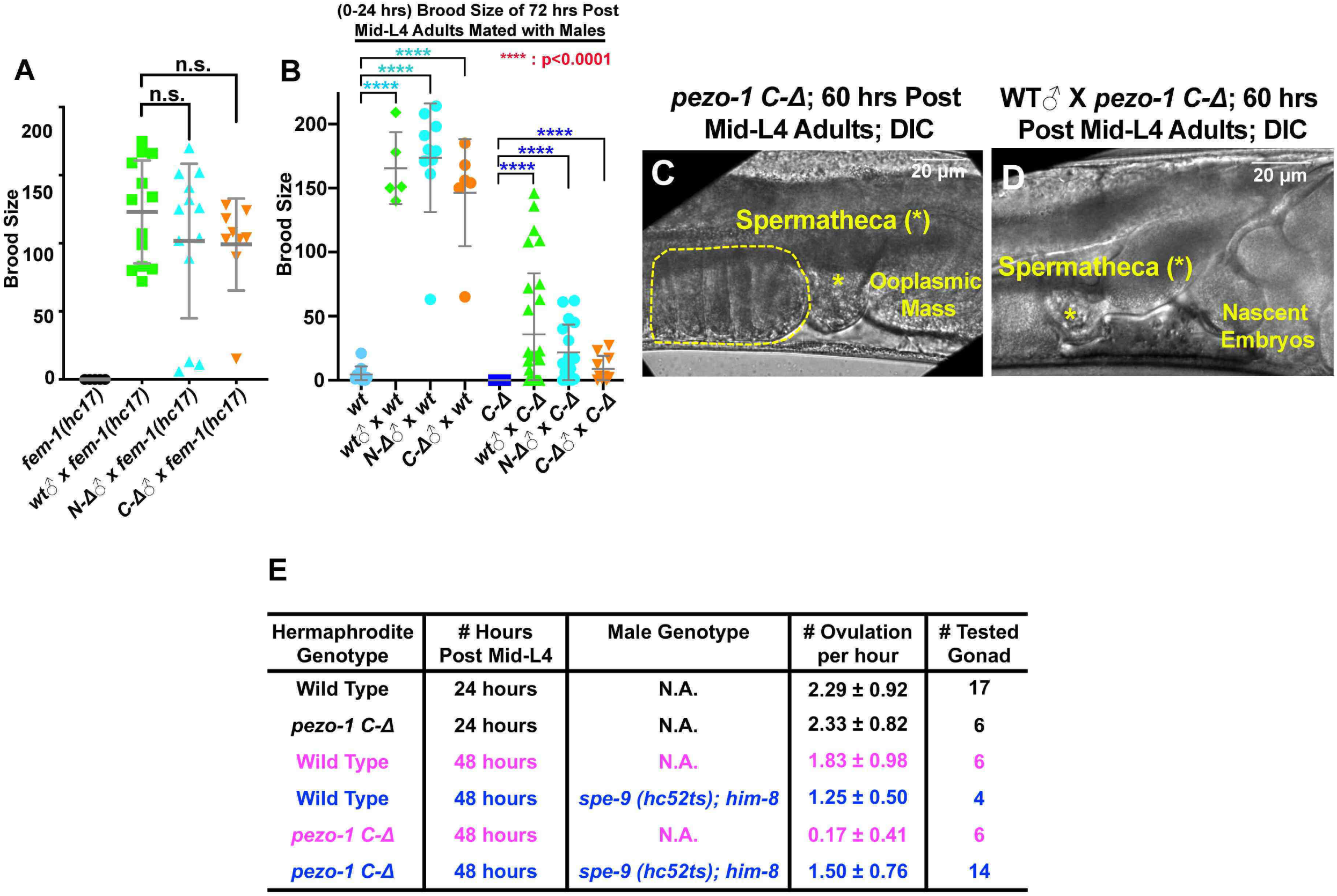
Male sperm rescue the ovulation defects in *pezo-1* mutants. (A) Both *pezo-1 C-Δ* and *N-Δ* males are fertile and sire progeny when mated with *fem-1(hc17ts)* mutants (essentially female animals). (B) Mating with male sperm rescued fertility in day 3 *pezo-1 C-Δ* adults (72 hours post mid-L4). (C) The oocyte maturation and ovulation rate are very low in day 3 *pezo-1 C-Δ* mutant adults and oocytes accumulate in the proximal gonad arm (yellow dashed circle). (D) In contrast, the ovulation rates are recovered to high levels after mating with wildtype male sperm. Newly-fertilized embryos pushed the ooplasmic mass out of the uterus. Yellow asterisk indicates the spermatheca (C, D). (E) Quantification of the oocyte ovulation rate of wildtype and *pezo-1 C-Δ* adults at different ages. *spe-9 (hc52ts)* sperm significantly rescue ovulation rates in *pezo-1 C-Δ* hermaphrodites even though they do not fertilize oocytes. P-values: **** <0.0001 (*t*-test). Scale bars are indicated in each panel.

### Sperm guidance and navigation is disrupted in *pezo-1* mutants

In wildtype hermaphrodites, the sperm are constantly being pushed out of the spermatheca each time the sp-ut valve opens to expel the fertilized oocyte into the uterus. These sperm, however, are fully capable of crawling back to the spermatheca to induce high levels of oocyte maturation and ovulation (Miller, 2001; Miller et al., 2003). This is a very efficient mechanism such that almost every sperm in a hermaphrodite is used to fertilize an oocyte. It is sperm number that defines brood size; oocytes are in abundance. Oocytes secrete F-series prostaglandins derived from polyunsaturated fatty acids (PUFAs) to guide sperm to the spermatheca (Han et al., 2010; Kubagawa et al., 2006). To test whether *pezo-1* mutants fail to attract the sperm back to the spermatheca, male sperm navigational performance was assessed *in vivo* by staining males with a vital fluorescent dye, MitoTracker CMXRos, which efficiently stains sperm in live animals (Whitten and Miller, 2007). Both wildtype and *pezo-1 C-Δ* stained males were mated to non-labeled wildtype hermaphrodites for 30 minutes. The sperm distribution was assessed and quantified by dividing the uterus into three zones (Fig. 7A) and counting the number of fluorescent sperm in each zone (McKnight et al., 2014) one hour after males were removed from the mating plates. In wildtype hermaphrodites, most fluorescent sperm from both wildtype and *pezo-1 C-Δ* navigated through the uterus and accumulated in the spermatheca (Fig. 7B, D, F and H). Similar observations were also found when stained wildtype and *pezo-1 C-Δ* sperm were tested in the *fem-1* hermaphrodites (data not shown). However, fewer sperm reached the spermatheca in Day-3 adult *pezo-1 C-Δ* hermaphrodites and most sperm remained throughout zone1 and zone 2, the zones furthest from the spermatheca (Fig. 7C, E, G, and H). This was observed for both wildtype and *pezo-1* mutant male sperm in mating with *pezo-1 C-Δ* hermaphrodites (Fig. 7H). These observations suggest that in the wildtype hermaphrodite reproductive tracts, *pezo-1* mutant male sperm are motile and display normal navigational behavior. However, in *pezo-1* mutant hermaphrodite reproductive tracts, both wildtype and *pezo-*1 mutant sperm were compromised in their navigational behavior over the time frame of this experiment. Alternatively, the ooplasmic masses that accumulate in the uterus from the crushed oocytes in *pezo-1* mutants could be interfering with the migration of wildtype and *pezo-1* mutant sperm back to the spermatheca.

**Figure 7.**
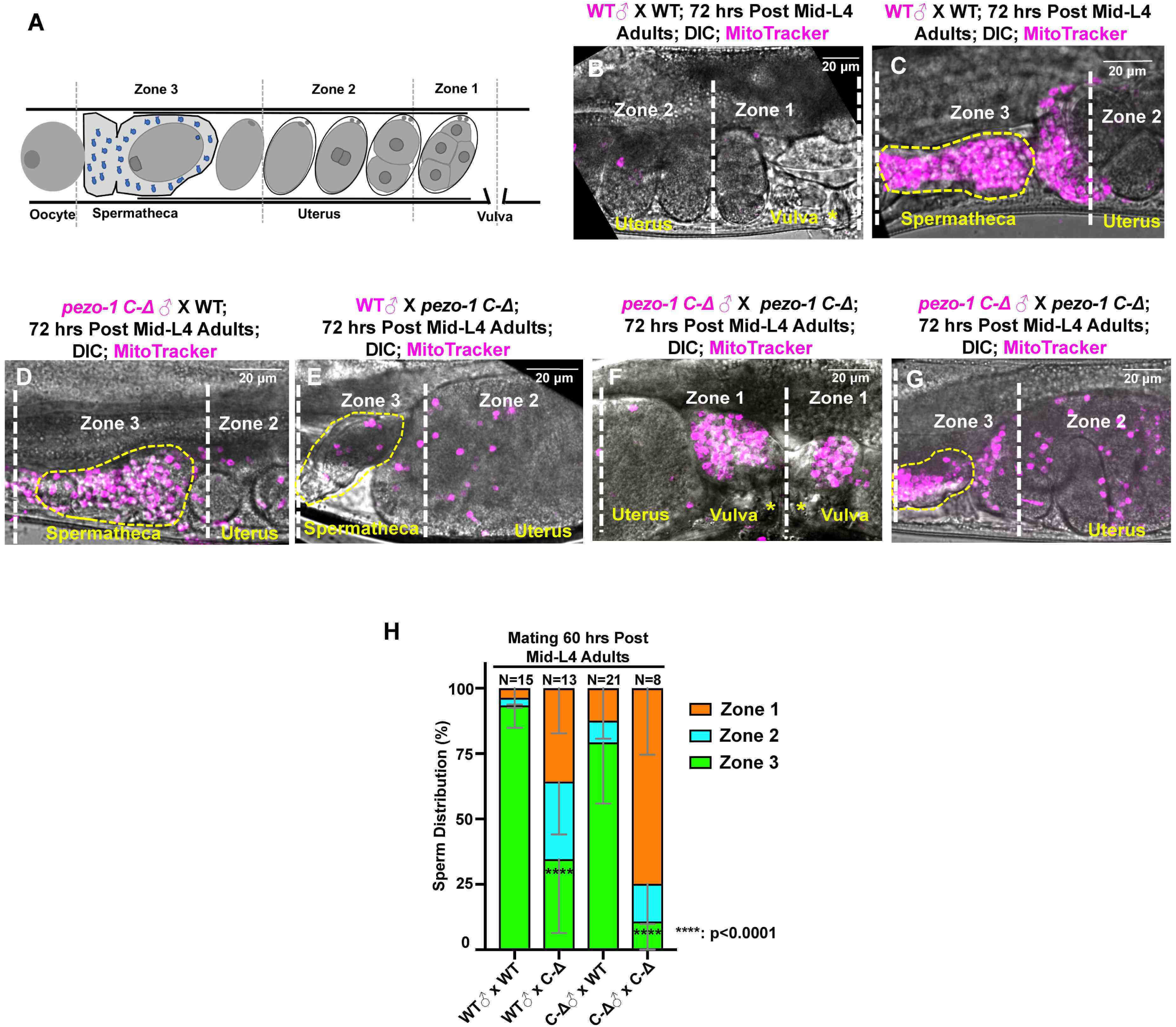
Sperm guidance and navigation is disrupted in *pezo-1* mutants. (A) To quantify sperm migration, this illustration indicates the three zones that were scored for sperm distribution. Zone 3 is the spermatheca (yellow dotted circles in panels C-E, G) while Zone 1 is the area closest to the vulva. Sperm distribution is measured 1 hour after males were removed from the mating plate. (B-G) The distribution of fluorescent male sperm labeled with MitoTracker in the three zones in both wildtype and *pezo-1* mutants 1 hour after the males were removed. (H) Quantification of sperm distribution values. Number of the scored uteri is shown above each of the bars. P-values: **** <0.0001 (*t*-test). Yellow asterisks indicate the vulva (F). Scale bars are indicated in each panel.

### Tissue-specific degradation of PEZO-1 reveals multiple roles of PEZO-1 in both somatic tissues and germline cells

Our study aims to understand the role of PEZO-1 in regulating reproduction and coordinating inter-tissue signaling. To dissect PEZO-1 function in distinct tissues, we utilized an auxin-inducible degradation system (AID) to degrade PEZO-1 in the soma and the germ line (Zhang et al., 2015). We knocked-in the degron coding sequence at the *pezo-1* C-terminus using CRISPR/Cas 9 so that all isoforms would be targeted (Fig. 8A). To activate the AID system, this line was then crossed with the strains expressing the degron interactor transgene *tir-1::mRuby* driven by the following promoters: *Peft-3, Ppie-1* and *Psun-1*. *Peft-3::tir-1::mRuby* was expressed in most or all somatic tissues, including the spermatheca and the sheath cells (Fig. 8B) while *Ppie-1::tir-1::mRuby* and *Psun-1::tir-1::mRuby* were expressed in the germ line (Fig. 8C, D). Weak TIR1-1::mRuby expression was observed in the sperm of the germline strains (Fig. 8C, D). To assess the defects associated with the degradation of PEZO-1 in these different tissues, L4 animals were exposed to either 0.25% ethanol as control or 1-2 mM auxin (indole-3-acetic acid, or IAA) and brood sizes were determined 0-60 hours post L4 (Day 1-3). Interestingly, the brood sizes were significantly reduced in each of the PEZO-1::Degron strains compared with control, regardless of the promoter used. However, the reduction in brood size was less severe than observed in the *pezo-1^ko^* mutants (Fig. 8E, F, 2A). To ensure efficient degradation, we exposed animals to auxin for one generation and analyzed the brood size in their F1 progeny. This longer auxin exposure did not significantly enhance the reduction in brood size (data not shown).

**Figure 8.**
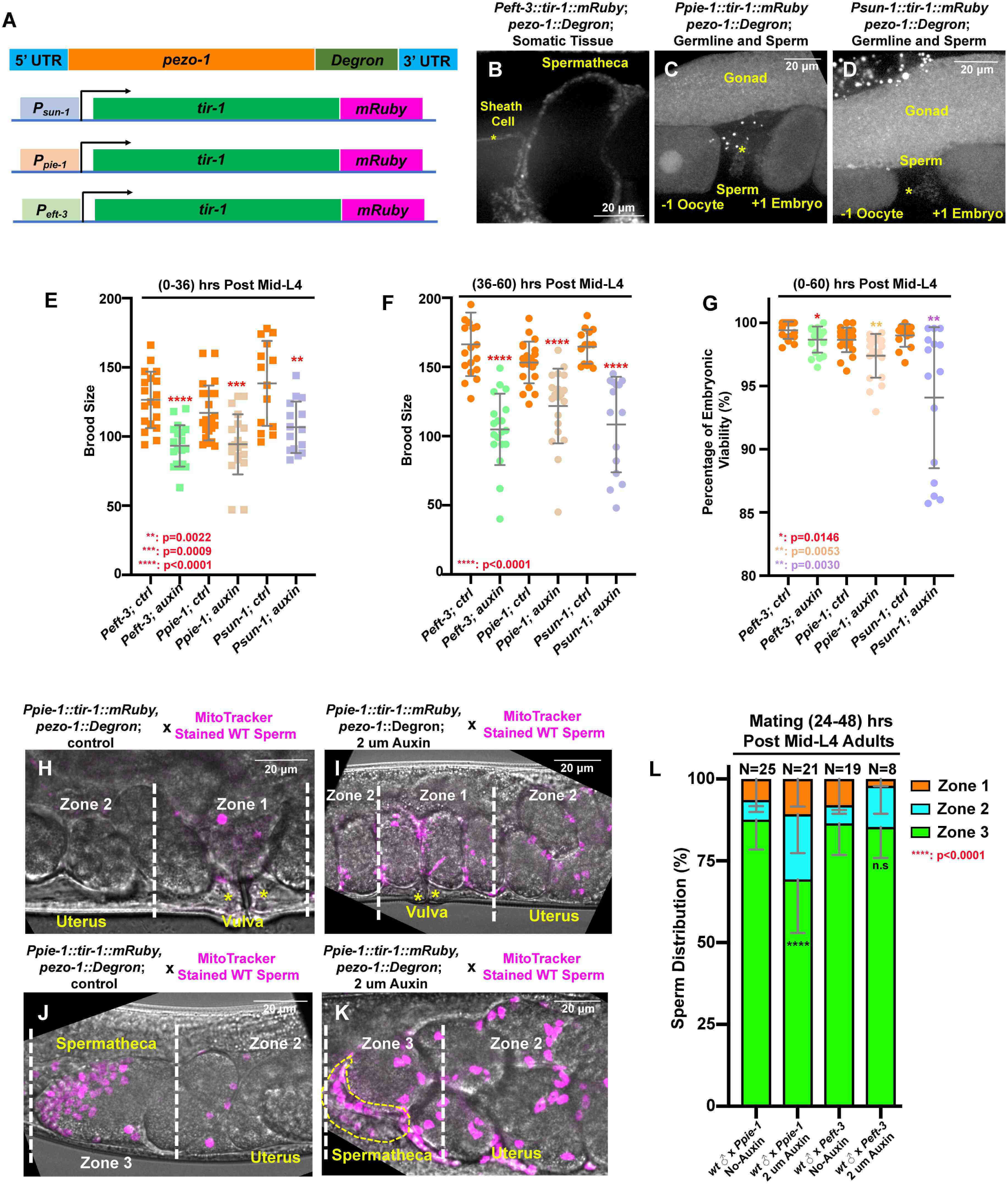
Tissue-specific degradation of PEZO-1 displays a reduced brood size and causes sperm navigational defects. (A) Schematic of the auxin-inducible degradation (AID) system. A degron tag was inserted at the 3’ end of the *pezo-1* coding sequence using CRISPR/Cas9-mediated editing. (A-C) The *eft-3* promoter was used to drive TIR-1 expression in most or all somatic tissues, including the spermatheca and the sheath cells (A, B). TIR-1::mRuby driven by the germline specific promoters, *sun-1* and *pie-1*, is strongly expressed in the germline and oocytes (A, C, D). Weak expression of TIR-1::mRuby was also observed in the sperm (C, D, asterisk). (E-G) Brood size and embryonic viability were reduced in all degron strains when animals were treated with 1 mM or 2 mM auxin. Data are presented as the mean ± standard error from at least two independent experiments. (H-K) Sperm distribution 1 hour after male removal from mating plates. The germline specific PEZO-1::Degron hermaphrodites were mated with wildtype males for 30 minutes. The representative images show that *pezo-1* degradation in the germ line influences sperm distribution from the vulva (zone 1) to the spermatheca (zone 3). (L) Quantification of sperm distribution in the PEZO-1::Degron strain grown on plates with (+) or without (−) 2 mM auxin. P-values: * = 0.0146 (G); ** = 0.0022 (E); ** = 0.0030 (G); ** = 0.0053 (G); *** = 0.0009; **** <0.0001 (E, F, L) (*t*-test). Scale bars are indicated in each panel.

Depletion of PEZO-1 in the somatic tissues, including spermathecal and sheath cells, led to a variety of ovulation defects (Fig. S4A-I). Pinched oocytes were frequently observed during ovulation (N= 9/27, Fig. S4I). A fraction of the pinched oocytes entered the spermatheca, while the rest were left in the oviduct (Fig. S4C, D, I). Surprisingly, most of the pinched oocytes were successfully expelled into the uterus and underwent embryogenesis as smaller embryos (data not shown). Additionally, the process of oocyte entry into the spermatheca was frequently delayed or blocked (Fig. S4E-I), suggesting the distal spermathecal valve remained closed. In experiments in which wildtype sperm were *in vivo* labeled as described above, and mated into control and somatic-specific PEZO-1::degron hermaphrodites, nearly 90% of the labeled sperm reached the spermatheca (zone 3) and only a few labeled sperm were observed in the uterus (Fig. 8H, J, L). Notably, the ooplasmic uterine masses that we observed in our *pezo-1^ko^* mutants were rarely observed in the somatic-specific degron strain.

Consistent with our male mating experiments, only 69% of the MitoTracker-labelled wildtype sperm accumulated at the spermatheca (zone 3) in the germline-specific PEZO-1::degron animals exposed to the auxin (Fig. 8H-L). The remaining sperm were observed throughout the whole uterus (zone 1 and zone 2) after one hour of mating (Fig. 8I, K, L). Crushed oocytes were rarely observed in the uterus of the germline-specific PEZO-1::degron animals, in which the sperm distribution assay was performed. Therefore, degradation of PEZO-1 in the germ line did not cause the severe uterine ooplasmic masses as we have observed for our *pezo-1^ko^* mutants but did interfere with sperm navigation to the spermatheca, likely due to the impaired attractant signaling. This is a more likely explanation since uterine ooplasmic masses are not a physical impediment to account for the defects in sperm migration.

### Disrupting PEZO-1 reveals multiple roles in inter-tissue signaling

Given that PEZO-1 expresses in all reproductive tissues and oocyte transit defects are striking in our *pezo-1* mutants, we reasoned that *pezo-1* may function in the communication between sheath cells and germ cells. The innexin gap junction genes *inx-14* and *inx-*22 play a role in the communication of gonadal sheath cells with germ cells as well as in the attraction of sperm back to the spermatheca (Edmonds et al., 2011). RNAi of either *inx-14* or *inx-*22 enhanced the low brood size phenotype of the *pezo-*1 mutants (Fig. S5A, B). In *pezo-1* mutants, more than an additional 50% reduction in brood size was observed after *inx-14* and *inx-22* RNAi (Fig. S5A, B). At least 30% of the *pezo-1* mutant embryos displayed abnormal shape and failed to hatch after *inx-14* RNAi treatment (Fig. S5A, B). The enhancement of phenotypes with these innexins suggests that PEZO-1 may be involved in regulating inter-tissue signaling.

Intestinal yolk granules containing PUFA precursors are endocytosed by oocytes and are used by the oocytes to synthesize F-series prostaglandins to guide sperm to the spermatheca (Han et al., 2010; Kubagawa et al., 2006). These yolk granules can be visualized by a yolk protein YP170::tdimer2 transgene (Fig. S6A, A’). Unlike wildtype animals, yolk accumulation was observed in the pseudocoelomic cavity surrounding the gonad of the *pezo-1* mutants (Fig. S6B, C, B’, C’), suggesting that yolk endocytosis into the oocytes was defective. The reduced endocytosis of yolk may disrupt prostaglandin synthesis in the oocyte, which may lead to a defect in the oocytes’ ability to attract sperm towards the spermatheca. As shown earlier, the proper number of sperm are made in our *pezo-1* mutants but they fail to navigate back to the spermatheca after being washed out of the spermatheca by the process of ovulation and fertilization. Collectively, PEZO-1 may serve as an important signaling regulator between sheath cells, oocytes, and sperm, the detailed mechanism of which will be the focus of future studies.

### Modeling human PIEZO genetic diseases in *C. elegans*

*PIEZO* patient-specific alleles, which are known to disrupt the normal physiological functioning of the cardiovascular, musculoskeletal, and blood systems in humans, were the motivation for examining the role of *pezo-1* in the tubular structures of *C. elegans.* Our studies with null alleles of *pezo-1* provide strong evidence that *pezo-1* is essential for *C. elegans* reproduction. It is therefore reasonable to model human monogenic diseases associated with *PIEZO1 and PIEZO2* mutations using the *C. elegans* reproductive system as a read-out of function. Three missense mutations have been identified at a conserved arginine residue (p.R2488Q) in *PIEZO1* and (p.R2718L/P) in *PIEZO2* individuals; these individuals have been diagnosed with Dehydrated Hereditary Stomatocytosis (DHSt) and Distal Arthrogryposis type 5 (DA5), respectively (Andolfo et al., 2013; Coste et al., 2013; Li et al., 2018; McMillin et al., 2014). Previous studies have shown that these arginine changes are functioning as gain-of-function mutations in their respective PIEZO protein (Albuisson et al., 2013; Coste et al., 2013; Li et al., 2018; McMillin et al., 2014). Sequence alignment indicated that R2405 in *C. elegans* PEZO-1 is the homologous arginine residue to both R2488 in human PEIZO1 and R2718 in human PIEZO2 (Fig. 9A). Using CRISPR/Cas9, we generated the patient-specific *PIEZO2* allele (p.R2718P) in *C. elegans*, named *pezo-1 (R2405P)*. To compare this patient-specific allele with that of our null alleles, and to determine the phenotypic consequences of a patient-specific allele, homozygous animals carrying the *pezo-1 (R2405P)* mutation were created. Such homozygotes displayed reproductive defects similar to the *pezo-1^ko^* mutants, including reduced ovulation rates, ooplasmic uterine masses (Fig. 9B), and reduced brood sizes (Fig. 9C). Additionally, *pezo-1(R2405P)* homozygotes had genetic interactions with *itr-1* RNAi and *lfe-2* RNAi consistent with our findings with *pezo-1^ko^* mutants (Fig. 9D). Interestingly, similar to the rescue assay in *pezo-1 C-Δ*, the reduced ovulation rate in *pezo-1(R2405P)* was also significantly rescued by *spe-9(hc52ts)* sperm, suggesting that this variant of *pezo-1* may similarly disrupt ovulation and sperm signaling, leading to self-sterility (Fig. 9E). Overall, these observations support the idea that *C. elegans* is an appropriate model system to study *PIEZO* diseases. Future suppressor screens with this and other *pezo-1* patient-specific alleles should help identify other genetic interactors.

**Figure 9.**
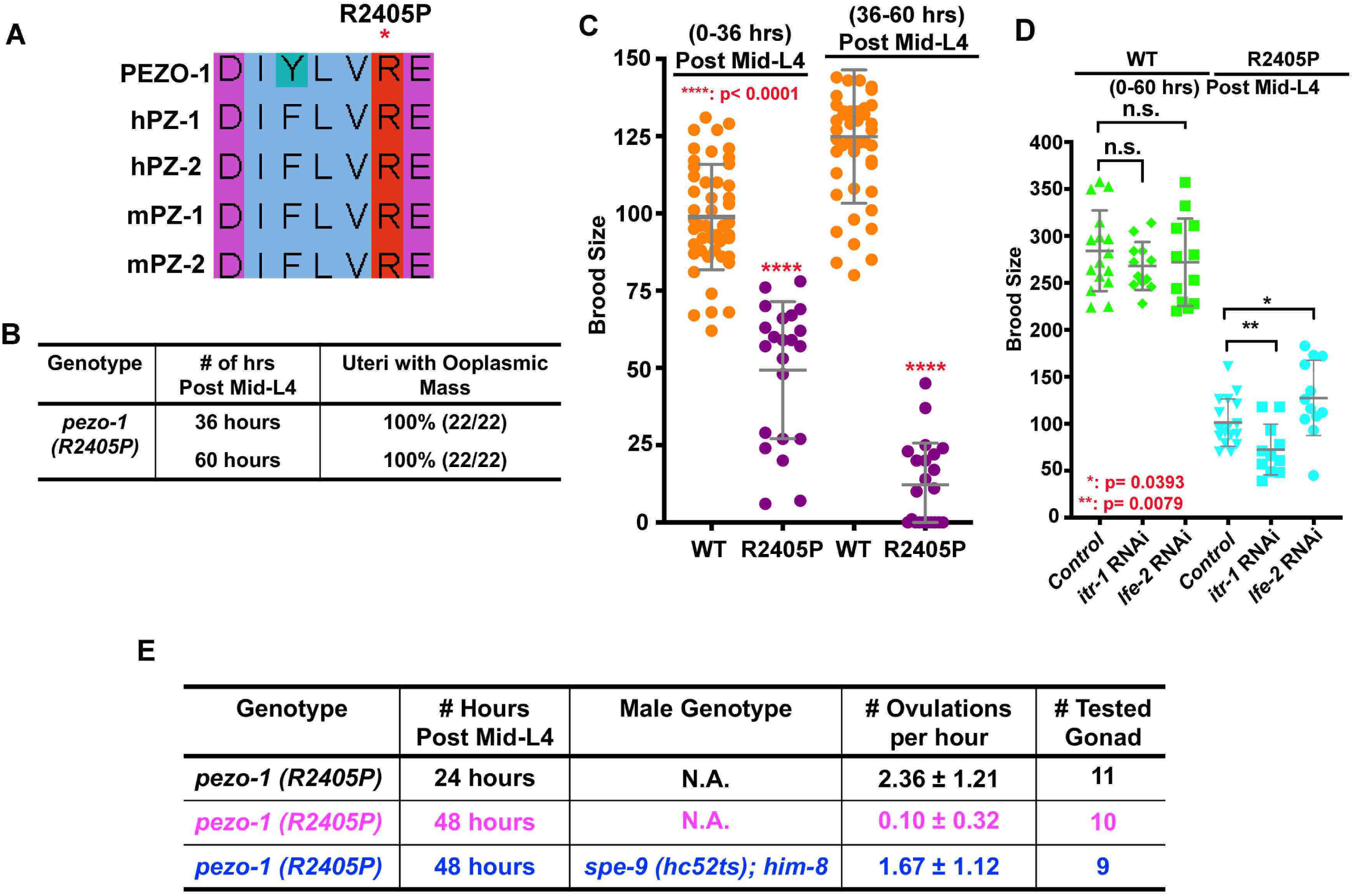
A *PIEZO1* disease allele causes severe brood size reduction in *C. elegans*. (A) Sequence alignment showing arginine 2405 (R2405) in *C. elegans* PEZO-1 is highly conserved with human and mouse PIEZO1 and PIEZO2. (B) A conserved patient-specific allele *pezo-1(R2405P)* was generated and causes uterine ooplasmic masses (B) and a severe reduction in brood size (C). (D) *itr-1 (RNAi)* enhanced the brood size reduction of *pezo-1 (R2405P)* mutants, while *lfe-2 (RNAi)* slightly rescued the reduced brood size. (E) *spe-9 (hc52ts)* sperm rescued the very low ovulation rate in *pezo-1 (R2405P)* hermaphrodites. P-values: * = 0.0393 (D); ** = 0.0079 (D); **** <0.0001 (C) (*t*-test).

## Discussion

The PIEZO proteins are pore-forming protein complexes, responsible for sensing mechanical stimuli during physiological processes. Most studies of PIEZOs have focused on electrophysiological assays in cultured cells. To take advantage of an *in vivo* system to investigate the developmental roles of the PIEZO channel in mechanotransduction, we have generated deletion as well as patient-specific alleles in the sole *C. elegans pezo-1* gene. The *C. elegans* reproductive system is an attractive tubular system to study *PIEZO* function and mimic the *PIEZO* patient-specific alleles, which are known to disrupt the normal physiological functioning of the cardiovascular, musculoskeletal, and blood systems in humans (Albuisson et al., 2013; Alper, 2017; Andolfo et al., 2013; Bae et al., 2013). Though the PEZO-1 protein is broadly expressed throughout the animal, we focused on the reproductive system given its striking phenotypes. Utilizing different *pezo-1* mutants and the tissue-specific degradation of PEZO-1, our data indicate that dysfunction of *pezo-1* led to a significantly reduced brood size. This reduced brood size phenotype worsens with age, suggesting perhaps that maternal PEZO-1 perdures enough to allow normal ovulations through the first day of adulthood. Alternatively, there is a potential parallel pathway that complements the loss of PEZO-1 in early adulthood that does not persist as the adult ages. In *C. elegans*, the complete reproductive process incorporates a series of sequential events, including proper ovulation, fertilization, expulsion of the fertilized zygote into the uterus and sperm navigation back to the spermatheca after each fertilization event, all of which are regulated by multiple inter-tissue signaling pathways.

### PEZO-1 channel regulates ovulation and expulsion of the fertilized zygote possibly through maintaining cytosolic Ca^2+^ homeostasis

Ovulation is driven by the rhythmic and coordinated contraction of the gonadal sheath cells and opening of the spermathecal distal valve (McCarter et al., 1999). Similarly, expulsion of the fertilized zygote into the uterus is achieved by the contraction of the spermatheca and opening of the spermathecal-uterine valve. Mutations in the *pezo-1* gene cause quite dramatic effects on this entire process. We observed sheath cell defects such that the mature oocyte was not properly pushed into the spermatheca. In addition, spermathecal valve defects either inhibited proper entry of the oocyte into the spermatheca, or proper exit. In many cases, the oocyte was crushed as it progressed through the spermatheca. The result often was the presence of an ooplasmic mass in the uterus. Contractility in both the sheath cells and the spermathecal cells are regulated by the IP3 receptor *itr-1* and (IP3) kinase *lfe-2* (Bui and Sternberg, 2002; Kovacevic et al., 2013). Genetic interactions between *pezo-1* mutants and *itr-1* or *lfe-2* RNAi support the idea that *pezo-1* is necessary for maintaining Ca^2+^ homeostasis during ovulation and zygote expulsion. This is consistent with previous studies showing PIEZO1 responses to mechanical stimuli through Ca^2+^ signaling (He et al., 2018; Li et al., 2014). Based on present studies, we hypothesize a few possible pathways for a Ca^2+^ mediated response to mechanical stimuli to which PEZO-1 may contribute. One possibility is that PEZO-1 may detect when cytosolic Ca^2+^ levels are extremely low and replenish the cell with extracellular Ca^2+^, in a similar manner as the CRAC channel ORAI-1 (Lorin-Nebel et al., 2007). Consistent with this idea, our genetic data revealed an enhancement of the *pezo-1* phenotype upon CRAC channel *orai-1* RNAi, which is responsible for replenishing cytosolic Ca^2+^ (Fig. 5C). This suggests that PEZO-1 and ORAI-1 act in parallel pathways to replenish cytosolic Ca^2+^. Previous studies identified the ER Ca^2+^ pump sarco/endoplasmic reticulum Ca^2+^ ATPase (SERCA) as an interacting partner of PIEZO1, which suppresses PIEZO1 activation (Zhang et al., 2017). SERCA is essential for recycling Ca^2+^ into SR/ER Ca^2+^ stores, which is an important process for maintaining Ca^2+^ homeostasis during tissue contractility (Periasamy and Huke, 2001; Zwaal et al., 2001). PIEZO1 has been reported to be involved in integrin activation to recruit the small GTPase R-Ras to the ER, which promotes Ca^2+^ release from an intracellular store to the cytosol (McHugh et al., 2010). Combined with the expression pattern of PEZO-1 at the ER region, these observations suggest that PEZO-1 may act as an ER Ca^2+^ channel to regulate ER Ca^2+^ homeostasis. Lastly, normal spermathecal GCaMP were observed during the first three ovulations in *pezo-1* mutants, suggesting that other Ca^2+^ or mechanosensitive channels may perform redundant functions during Ca^2+^ influx. One alternative model could be that PEZO-1 acts in parallel to these Ca^2+^ regulators and yet does not have a direct role in calcium homeostasis itself. Future studies will be required to resolve the precise molecular effect of PEZO-1 on Ca^2+^ and understand how PEZO-1 regulates inter/intra cellular communication with/without Ca^2+^ and potentially how other interacting partners coordinate during these processes.

### PEZO-1 channel is required for sperm navigation and inter-tissue signaling

*C. elegans* employs multiple peptide and lipophilic hormones to coordinate different tissues during reproduction. Ovulation is initiated by MSP (major sperm proteins) signaling derived from sperm to trigger oocyte maturation and sheath cell contraction (Kuwabara, 2003; McCarter et al., 1999; Miller, 2001). After each fertilization event, oocytes secret F-series prostaglandins (F-PGs) into the extracellular environment of the reproductive tract and stimulate sperm attraction back to the spermatheca (Hoang et al., 2013). Our observations revealed a strong expression of PEZO-1 on both the plasma membranes of oocytes and sperm. Dysfunction of *pezo-1* causes a severe reduction of the ovulation rate and defective sperm navigation back to the spermatheca in the aged animals. Male mating significantly rescued the very low ovulation rate in *pezo-1* mutants. Furthermore, the sperm navigation defects were observed in the germline specific degradation of PEZO-1 animals, which showed less sperm successfully navigating back to the spermatheca. Collectively, depletion of PEZO-1 dsirupted the navigation of sperm to return to the spermatheca, which may lead to the reduced ovulation rate and defective sheath cell contraction.

The oocyte is rich in PUFAs that are mainly delivered to the oocytes via the intestine through yolk lipoprotein complex endocytosis. The oocyte converts the PUFAs into F-series prostaglandins (F-PGs), which it secretes to signal the sperm (Hoang et al., 2013; Kubagawa et al., 2006). Therefore, accumulation of yolk protein in the body cavity of *pezo-1* mutants may suggest a deficiency in yolk endocytosis (Fig. S6B-C’); this may have multiple consequences, one being that F-PGs synthesis in the oocyte is impaired. This impairment potentially leads to defects in the attractive signals that are necessary for sperm to navigate back into the spermatheca after being washed out by fertilization. At this time, however, it is unclear whether PEZO-1 is involved in the regulation of yolk granule endocytosis and subsequent prostaglandin synthesis.

Lastly, genetic interaction between *pezo-1* and the genes encoding the connexin hemichannel proteins, INX-14 and INX-22 (Fig. S5A, B), which function in oocyte maturation and sperm attraction (Edmonds et al., 2011), are consistent with the hypothesis that both *pezo-1* deletion and *inx-14/inx-22* RNAi inhibit oocyte-sperm communication (Edmonds et al., 2011; Govindan et al., 2009). Our study suggests that PEZO-1 may be involved in mediating the communication between oocyte and sperm to trigger ovulation as well as the navigational signaling to guide the sperm back to the spermatheca after each ovulation.

### Working Model

Our study supports the working model that PEZO-1 may function to promote the sheath cell contractions that push the oocyte into the spermatheca as the first step in ovulation (Fig. 10, step one). Simultaneously, PEZO-1 may sense the sheath cell contractions to open the spermathecal distal valve to allow oocyte entry into the spermatheca. During fertilization, the distal and spermathecal-uterine valves have to remain closed, which may also be governed by PEZO-1 (Fig. 10, step two). After fertilization, PEZO-1 may regulate the spermathecal tissues and control the sp-ut valve to trigger a series of events to expel the fertilized oocyte into the uterus. Lastly, PEZO-1 may function in the oocyte to attract the sperm back into the spermatheca after being pushed out by the exiting newly-fertilized oocyte (Fig. 10, step three). Thus, dysfunction of PEZO-1 may contribute to multiple defects in all these steps, including failure of oocyte entry into the spermatheca, the crushing of oocytes as they transit through the ovary and spermatheca, and defective signaling perturbing the sperm from crawling back into the spermatheca after each ovulation (Fig. 10). Future studies are underway to more precisely determine the PEZO-1 function in each tissue using even more cell-specific promoters in the AID degradation system.

**Figure 10.**
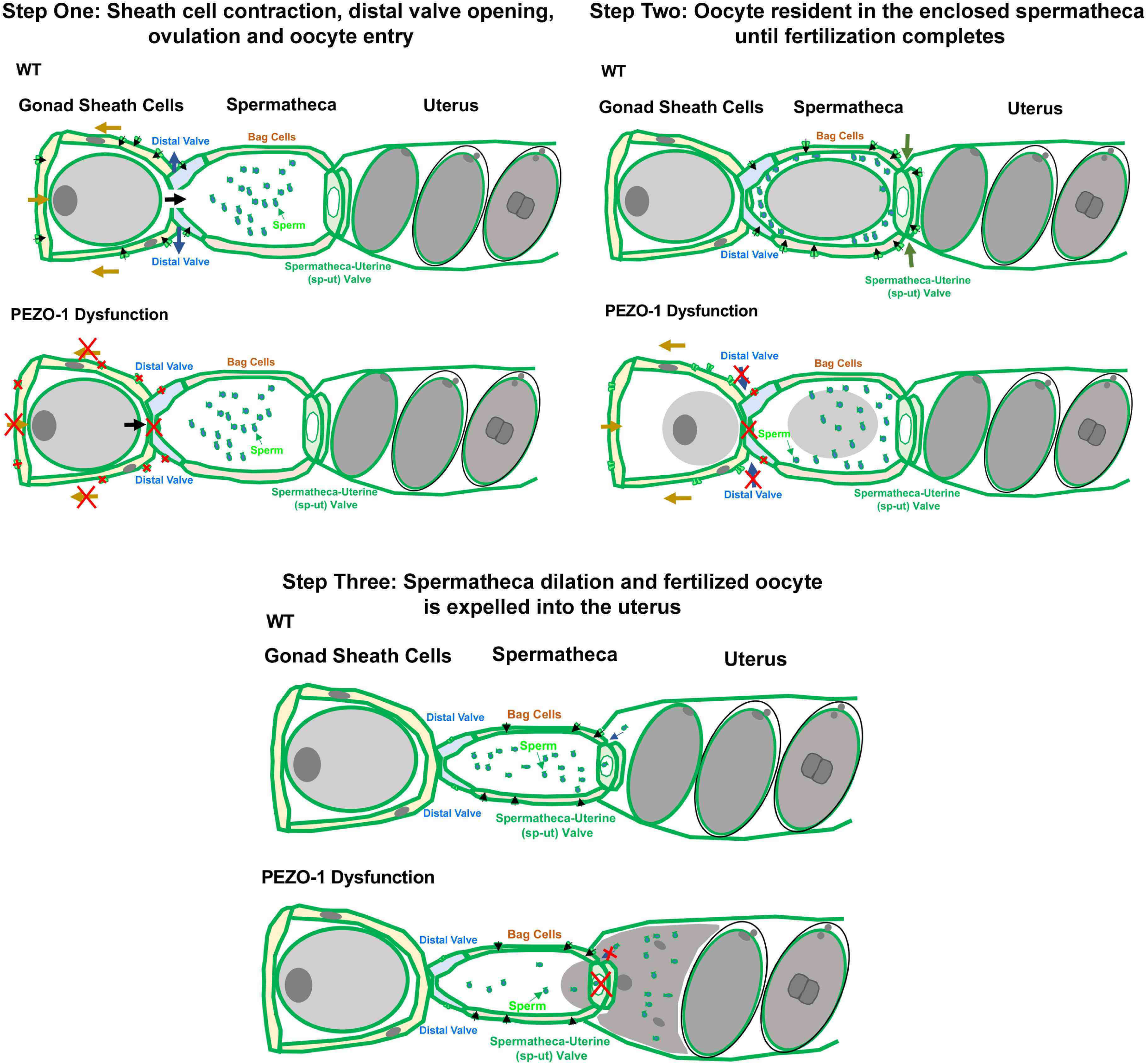
Working Model for PEZO-1 during ovulation. PEZO-1 regulates somatic sheath cells and the spermathecal distal valve to push the oocyte into the spermatheca (Steps One and Two). Dysfunction of PEZO-1 either blocks oocyte entry into the spermatheca (Step One) or pinches/crushes the oocytes when they transit from the ovary to the spermatheca (Step Two). During fertilization, the distal and spermathecal-uterine valves have to remain closed for 3-5 minutes (Step Two), which is governed by PEZO-1. Because of the earlier dysfunction of PEZO-1, only partial oocytes are enclosed in the spermatheca. After fertilization, PEZO-1 coordinates the spermathecal bag cells and the sp-ut valve to trigger a series of events to expel the oocyte into the uterus. Dysfunction of PEZO-1 crushes oocytes as they are expelled from the spermatheca (Step Three). PEZO-1 also functions to attract the sperm back to the spermatheca. Loss of PEZO-1 disrupts the navigation of the sperm from crawling back to the spermatheca.

### Modeling PIEZO diseases in the *C. elegans* reproductive system

Clinical reports indicate that either gain-of-function or loss-of-function mutations in the human *PIEZO1* and *PIEZO2* cause a variety of physiological disorders (Alper, 2017). Interestingly, both gain-of-function and loss-of-function missense mutations were identified in the same PIEZO disease, such as hydrops fetalis and lymphatic dysplasia. However, the molecular mechanism underlying both extremes of PIEZO channel dysfunction remains unclear (Alper, 2017). Complete knockout of *PIEZO1 and PIEZO2* in mammalian models results in embryonic lethality and fetal cardiac defects, suggesting the important role of PIEZO1/2 in embryonic and cardiac development (Ranade et al., 2014; Zhang et al., 2019). However, lack of surviving homozygous *PIEZO1/2* mutants in mammalian models make it challenging to investigate the PIEZO function during embryogenesis and development.

A DA5 patient-specific allele in the *C. elegans pezo-1* gene displayed identical reproductive phenotypes as our *pezo-1* deletion mutants, suggesting that this allele must be loss-of-function. The observation that our *pezo-1* deletion strains and a putative patient-specific gain-of-function mutation both lead to reproductive defects suggests that either hypomorphic or hypermorphic PEZO-1 channel activity is harmful. Therefore, our study demonstrates the usefulness of *C. elegans* as a model system to investigate PIEZO-derived human diseases. The dramatic reduction in brood size will allow us to screen plausible chemical antagonists and agonists for PIEZO1 and PIEZO2 patient-specific alleles *in vivo*. In summary, we have demonstrated that the *C. elegans PIEZO1/2* ortholog, *pezo-1*, is required for efficient reproduction, and demonstrate the utility of *C. elegans* for the study of PIEZO functions. Future studies will determine if other patient-specific alleles disrupt ovulation and sperm navigational signaling. The tissue-specific degradation system used in this report will also allow us to further dissect the responsible cells or tissues that influence each of the phenotypes we observed in this study. Future genetic and FDA-approved drugs screens will be used to identify putative suppressors in *pezo-1* mutants. These screens may provide insightful approaches for future clinical therapy.

## Materials and Methods

### *C. elegans* strains used in this study

*C. elegans* strains were maintained with standard protocols. Strain information is listed in Table 1. AG493, AG494 and AG495 were created by crossing AG487 (*pezo-1::degron*) males with hermaphrodites containing *ieSi65* [*sun-1p::tir1::sun-1 3’UTR + Cbr-unc-119(+)*] *II*, *ieSi57* [*eft-3p::tir1::mRuby::unc-54 3’UTR + Cbr-unc-119(+)*] *II, and fxIs1*[*Ppie-1::tir1::mRuby*] *I*, respectively. We screened the F3 adults for the presence of the TIR-1::mRuby transgene by microscopy and genotyped for the *pezo-1::degron* by PCR. AG532 was created by crossing *pezo-1(av146* [*gfp::pezo-1*]*) IV* males with the *unc-119(ed3); pwIs98* [*YP170::tdimer2 + unc-119(+)*] *III* hermaphrodites containing YP170::tdimer2. F3 adults with YP170::tdimer2 were genotyped by PCR screening for the *pezo-1^KO^* allele.

**Table 1.**
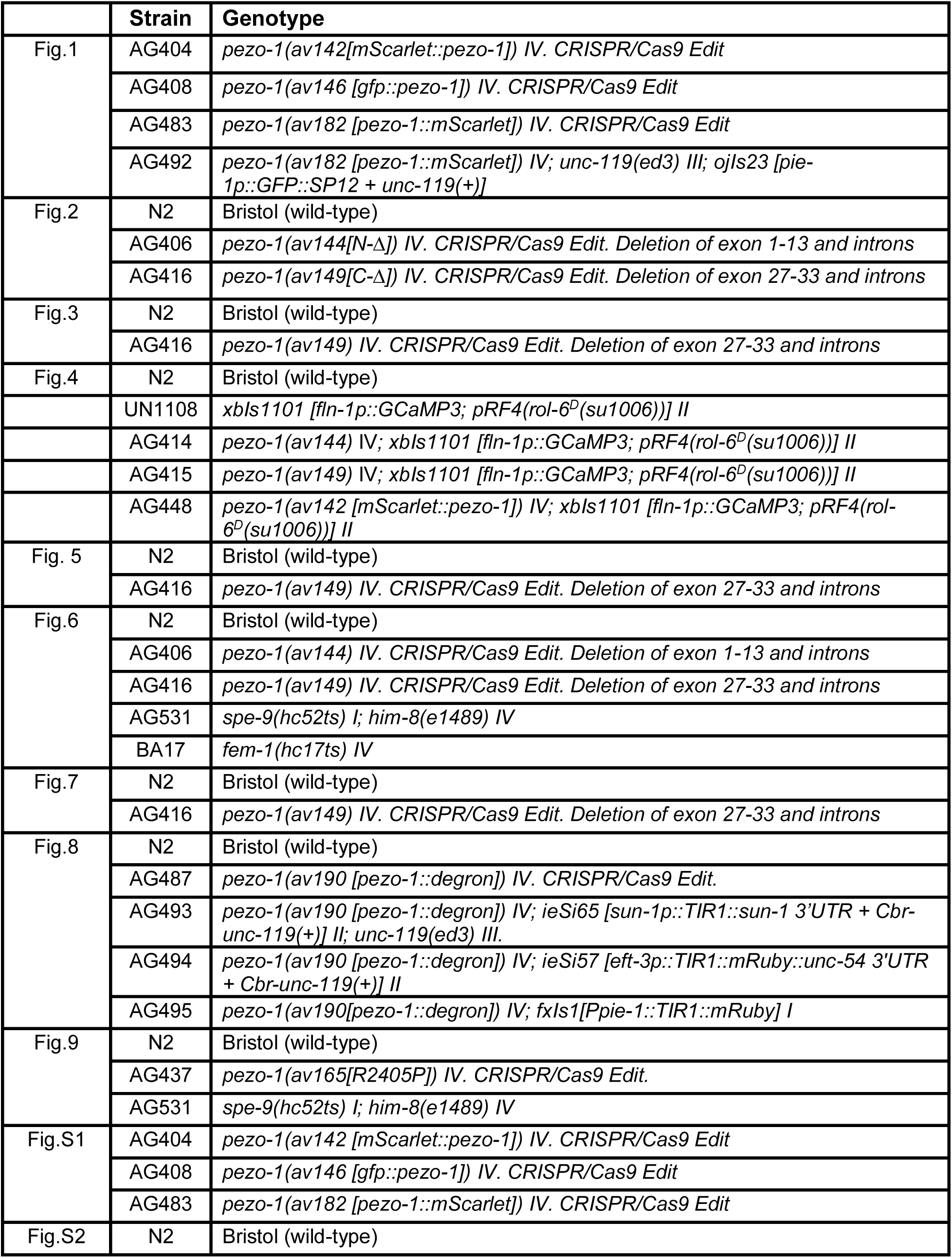
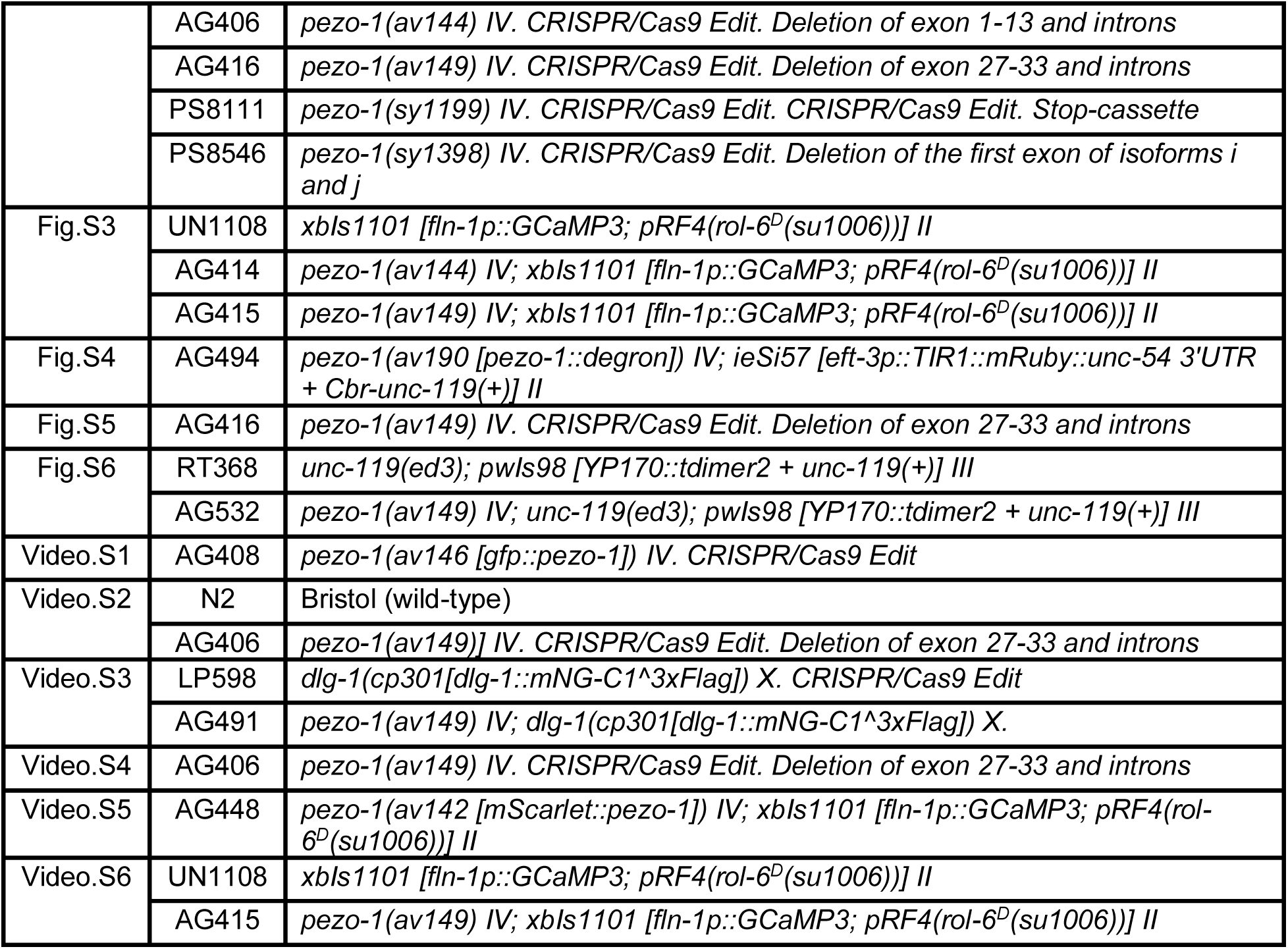
*C. elegans* strains list in the study.

### RNAi treatment

The RNAi feeding constructs were obtained from the Ahringer and Vidal libraries (Fraser et al., 2000; Rual et al., 2004). RNAi bacteria were grown until log phase was reached and spread on MYOB plates containing 1mM IPTG and 25 μg/ml carbenicillin and incubated overnight. To silence the target genes *itr-1*, *lfe-2*, *inx-14* and *inx-22*, mid-L4 hermaphrodites were picked onto plates with the IPTG-induced bacteria. Animals were grown on RNAi plates at 20°C for 36-60 hours. In order to improve the RNAi penetrance of *orai-1* and *sca-1*, L1 hermaphrodites were picked for RNAi feeding assays. Alternatively, mid-L4 hermaphrodites were incubated on the *orai-1* or *sca-1* RNAi plates for one generation, and F1 mid-L4 hermaphrodites were moved to fresh RNAi plates for brood size assays.

### Brood size determinations and embryonic viability assays

Single mid-L4 hermaphrodites were picked onto 35 mm MYOB plates seeded with 10 μl of OP50 bacteria and allowed to lay eggs for 36 hours (plate one contains the brood size from 0-36 hours post mid-L4). The same hermaphrodite was moved to a new 35 mm MYOB plate to lay eggs for another 24 hours and were removed from the plate (this plate contains the brood size from 36-60 hours post mid-L4). Twenty-four hours after removing the moms, only fertilized embryos and larvae were counted for the brood size. Brood sizes were determined at 36 hours and 60 hours. Percentage of embryonic viability= (the number of hatched larva / the total brood size) *100%.

### BODIPY 493/503 staining

BODIPY 493/503 (Invitrogen # D3922) was dissolved in 100% DMSO to 1 mg/ml. BODIPY stock was diluted by M9 to 6.7 μg/ml BODIPY (final concentration of DMSO was 0.8%) as the working stock. Hermaphrodites were washed in M9 three times and incubated in 6.7 μg/ml BODIPY for 20 minutes and washed again in M9 at least three times. All washes and incubations were performed in a concavity slide (ThermoFisher, # S99369). The stained hermaphrodites were anesthetized with 0.1% tricaine and 0.01% tetramisole in M9 buffer for 15-30 minutes. The anesthetized animals were then transferred to a 5% agarose pads for imaging. Image acquisition was captured by a Nikon 60 X 1.2 NA water objective with 1 μm z-step size.

### Whole animal DAPI staining

Animals were washed in M9 in a concavity slide, and then transferred to 1μl of egg white/M9/azide on SuperFrost slides (Daigger # EF15978Z). Alternatively, animals were directly picked from plates into egg white/M9/azide, trying not to carry over too much bacteria. With an eyelash, buffer around animals was spread out to a very thin layer, until animals are almost desiccated onto slide (too much egg white will blur final image and may give a halo when trying to observe DAPI). Slides were immersed in a coplin jar containing Carnoy’s fixative and fixed for a minimum of 1.5 hours and as long as one week at room temperature or 4°C. Sequential ethanol (EtOH) rehydration was carried out in coplin jars containing about 50 ml of the following solutions for 2 minutes each: 90% EtOH in water, 70% EtOH in water, 50% EtOH in PBS, 25% EtOH in PBS, and PBS alone. Slides were then immersed in coplin jars containing DAPI stain (1 μg/ml) in PBS for 10 minutes. Slides were rinsed three times, 5 minutes each, in PBS. A drop of Vectashield mounting medium (#H-1500-10) was added as was a coverslip, followed by nail polish to seal the coverslip. Image acquisition was captured by a Nikon 60 X 1.2 NA water objective with 1 μm z-step size.

### Live imaging to determine ovulation rates

For imaging ovulation, animals were immobilized on 4% agar pads with anesthetic (0.1% tricaine and 0.01% tetramisole in M9 buffer). DIC image acquisition was captured by a Nikon 60 X 1.2 NA water objective with 1-2 μm z-step size; 10-15 z planes were captured. Time interval for ovulation imaging is every 45-60 seconds, and duration of imaging is 60-90 minutes. Ovulation rate= (number of successfully ovulated oocytes) / total image duration.

### Imaging of yolk proteins in the animals

Day 1 adults (post mid-L4 24 hours) of both AG532 and RT368 were immobilized on 4% agar pads with anesthetic (0.1% tricaine and 0.01% tetramisole in M9 buffer). The acquisition of both DIC and 561 nm images were performed by our confocal imaging system (see below) with a Nikon 60 X 1.2 NA water objective. 20-30 z planes were captured with 1 μm z-step size. Images were generated by custom Fiji code using Image>Stacks>Z Project.

### CRISPR design

We used the Bristol N2 strain as the wild type for CRISPR/Cas9 editing. The gene-specific 20-nucleotide sequences for crRNA synthesis were selected with help of a guide RNA design checker from Integrated DNA Technologies (IDT) (https://www.idtdna.com) and were ordered as 20 nmol or 4 nmol products from Dharmacon (https://dharmacon.horizondiscovery.com), along with tracRNA. Repair template design followed the standard protocols (Paix et al., 2015; Vicencio et al., 2019). Approximately 30 young gravid animals were injected with the prepared CRISPR/Cas9 injection mix as described in the literature (Paix et al., 2015). *pezo-1 N-Δ* and *pezo-1 C-Δ* mutants were generated by CRISPR/Cas9 mixes that contained two guide RNAs at flanking regions of *pezo-1* coding regions. Heterozygous *pezo-1* deletion animals were first screened by PCR and then homozygosed in subsequent generations. mScarlet insertions at the *pezo-1* C-terminus were performed by Nested CRISPR (Vicencio et al., 2019). Homozygous *nest-1* edited animals were confirmed by PCR and restriction enzyme digestion and selected for the secondary CRISPR/Cas9 editing. Full-length mScarlet insertion animals were screened by PCR and visualized by fluorescence microscopy. All homozygous animals edited by CRISPR/Cas9 were confirmed by Sanger sequencing (Eurofins). The detailed sequence information of the repair template and guide RNAs are listed in Table 2.

**Table 2.**
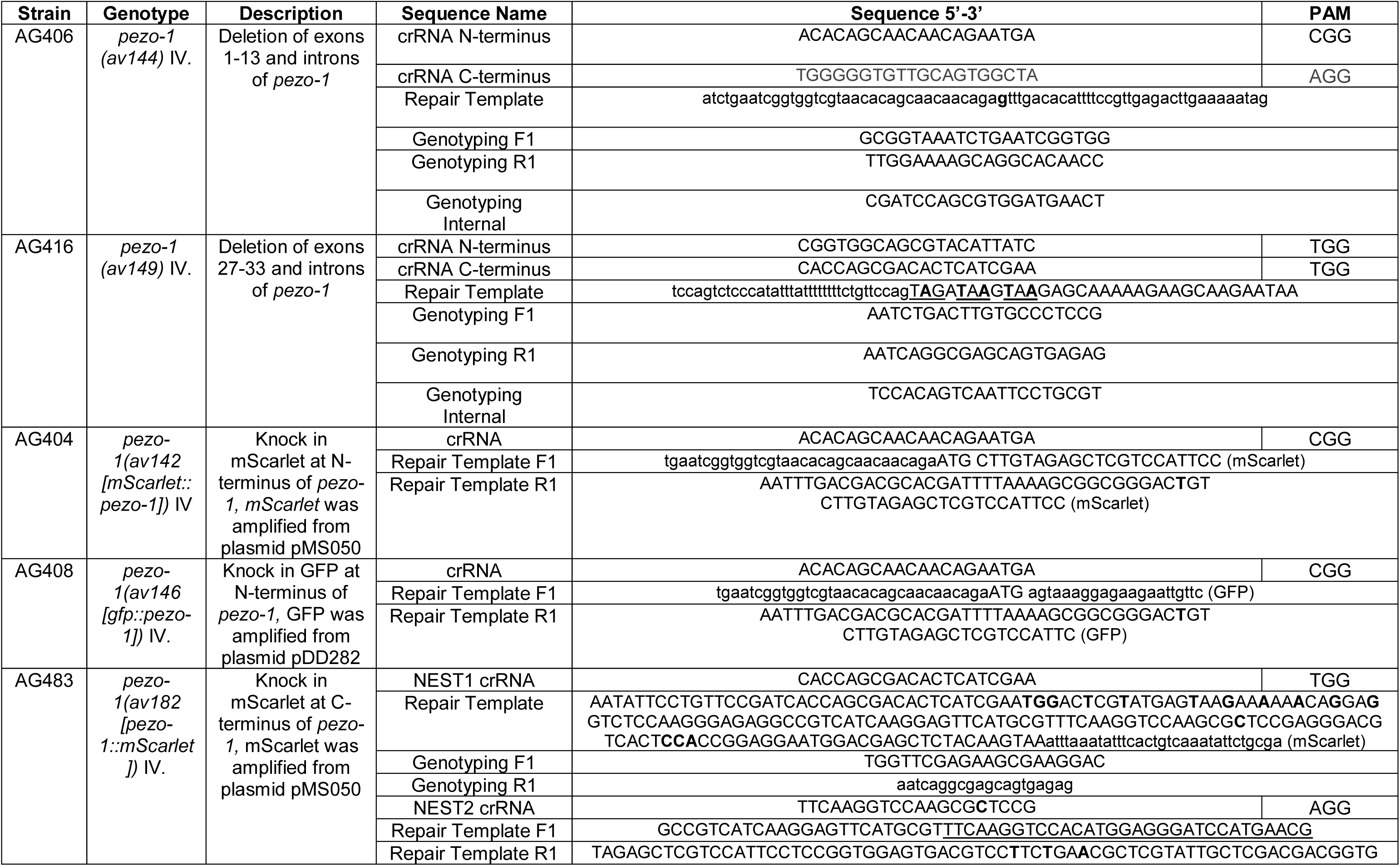
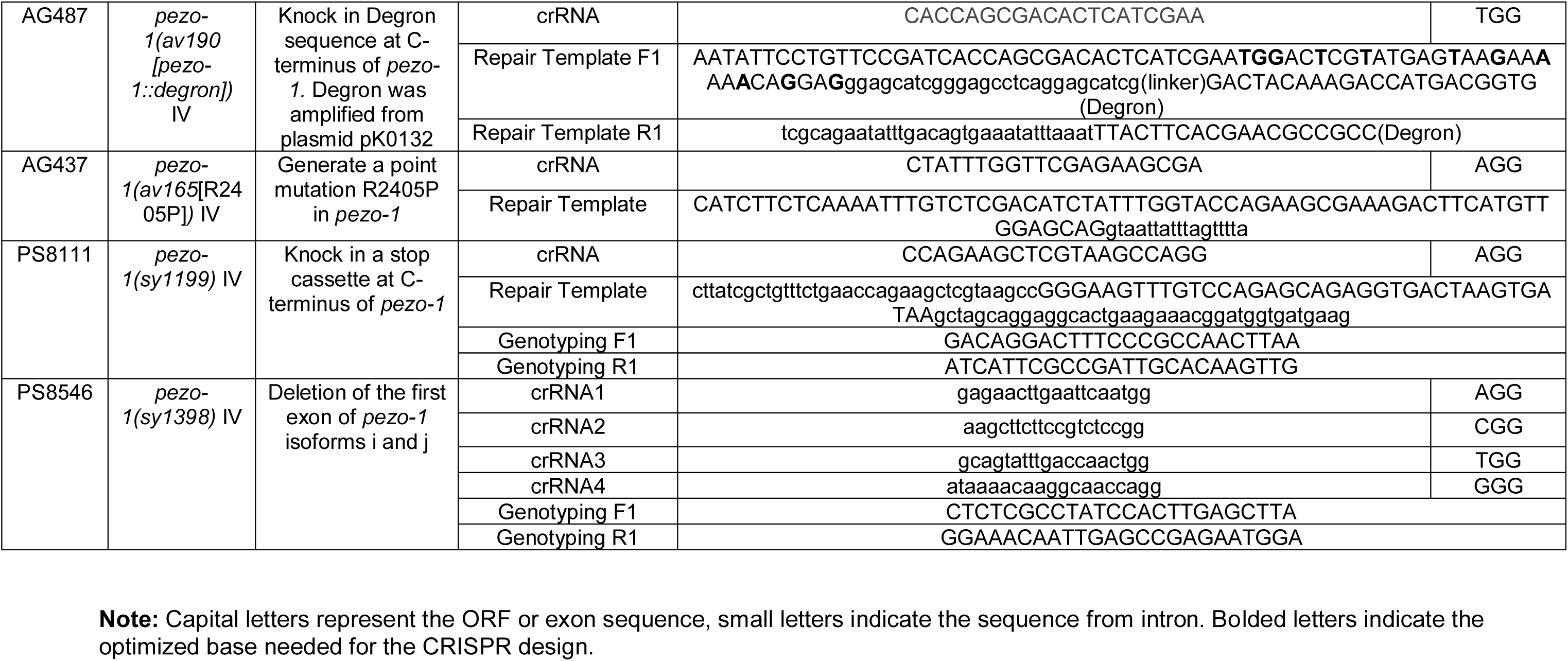
List of the sequence for the CRISPR design.

The short isoform deletion, *pezo-1 (sy1398)*, was generated using plasmid Cas9 expressed from a plasmid (Friedland et al., 2013) and four guides (GAGAACTTGAATTCAATGG, AAGCTTCTTCCGTCTCCGG, GCAGTATTTGACCAACTGG, ATAAAACAAGGCAACCAGG) along with a *dpy-10* guide and repair oligo. These reagents were injected into young adult N2 animals and successful injections were identified by the presence of roller or dumpy progeny on the plate. Roller progeny were singled out and screened via PCR for the deletion mutation. The deletion was verified by Sanger sequencing using two external primers (CTCTCGCCTATCCACTTGAGCTTA, GGAAACAATTGAGCCGAGAATGGA) to amplify the region. This deletion should only disrupt expression of isoforms I and j (Suppl. Fig. 2B). The CRISPR-Cas9 STOP-IN mutant, *pezo-1(sy1199)*, was generated using purified protein Cas9 at 10 μg/μl concentration, a purified guide RNA near the mutation location (CCAGAAGCTCGTAAGCCAGG), and a single stranded DNA repair oligo containing three stop codons, one in every reading frame (cttatcgctgtttctgaaccagaagctcgtaagccGGGAAGTTTGTCCAGAGCAGAGGTGACTAAGTGATAAgctagcaggaggcactgaagaaacggatggtgatgaag). These reagents were injected into N2 young adults along with a *dpy-10* guide and repair oligo. Successful injections were identified by the presence of dumpy and roller progeny. 30 roller progeny were singled out from ‘jackpot’ plates (plates with a high incidence of dumpy and roller progeny) and screened via PCR (GACAGGACTTTCCCGCCAACTTAA, ATCATTCGCCGATTGCACAAGTTG) and the presence of a NheI restriction site that was included in the repair oligo.

### Male mating assay with day 3 hermaphrodites

25-30 mid-L4 wildtype or *pezo-1* mutant hermaphrodites were isolated to a fresh growth plate for 60 hours (such animals should be day 3 adults at this time). To ensure mating success, ∼30 adult males and 10-15 day 3 hermaphrodites were transferred onto a 35 mm MYOB plate seeded with 10-20 μl of OP50 bacteria and allowed to mate for 12 hours. The other 10-15 day 3 hermaphrodites were singled and transferred to 35 mm MYOB plates seeded with 10 μl of OP50 bacteria as the controls. After the group mating, single mated hermaphrodites (72 hours post mid-L4) and 3-5 adult males were then transferred to a fresh 35 mm growth plate where mating could continue for another 24 hours. After 24 hours, the hermaphrodites (96 hours post mid-L4) and males were removed. The brood size (those embryos laid between 72-96 hours post mid-L4) and embryonic viability were determined 24 hours later after removal of all adults. Meanwhile, the broods from 60-96 hours post-mid L4 were also determined for the other 10-15 unmated day 3 hermaphrodites that were kept on single plates as controls.

### Mating assay with *fem-1* mutant

10-15 mid-L4 BA17 *fem-1(hc17ts)* hermaphrodites raised from embryos at the non-permissive temperature of 25°C were picked to mate with ∼30 adult males for 12 hours at 25°C. Single mated hermaphrodites and 3-5 males were then transferred to a fresh 35 mm growth plate and allowed to mate for another 24 hours at 25°C before all adults were removed from the plates. As control, 10-15 unmated BA17 hermaphrodites grown at 25°C were kept on single plates. The brood sizes and embryonic viability were determined 24 hours later. Alternatively, 10-15 L1 BA17 *fem-1(hc17ts)* hermaphrodites were isolated on a fresh growth plate and incubated at 25°C for 48 hours (young adult hermaphrodites). ∼30 adult males and 10-15 BA17 young hermaphrodites were then transferred onto a 35 mm MYOB plate seeded with 10-20 ul of OP50 bacteria and allowed to mate for 12 hours at 25°C. Single mated hermaphrodites and 3-5 males were then transferred to a fresh 35 mm growth plate. After laying embryos for 24 hours, the hermaphrodites and males were removed. Meanwhile, the other same age 10-15 unmated day 3 hermaphrodites were kept on single plates as the control. The brood size and embryonic viability were counted 24 hours later after removal of all adults. All the animals were incubated at 25°C during mating and propagation to ensure the penetration of the *fem-1(hc17ts)* phenotype.

### Mating assay with *spe-9* mutant

10-15 hermaphrodites were picked to mate with ∼30 AG521 [*spe-9(hc52ts)*] adult males for 12 hours at 25°C. Mated hermaphrodites were immobilized on 4% agar pads with anesthetic (0.1% tricaine and 0.01% tetramisole in M9 buffer) for ovulation rate assays. The acquisition of DIC images was performed by confocal imaging system (see below) with a Nikon 60 X 1.2 N with 1-2 μm z-step size and 10-15 z planes. Time interval for ovulation imaging is every 45-60 seconds, and duration of imaging is 60-90 minutes. Ovulation rate= (number of successfully ovulated oocytes) / total image duration.

### Sperm distribution assay and mating assay

MitoTracker Red CMXRos (MT) (Invitrogen # M7512) was used to label male sperm following the protocol adapted from previous studies (Hoang et al., 2013; Kubagawa et al., 2006). MT was dissolved in 100% DMSO to 1 mM. About 100 males were transferred to a concavity slide (ThermoFisher, # S99369) with 150 μl 10 μM MT solution (diluted in M9 buffer). Males were incubated in the MT buffer for 2 hours and then transferred to fresh growth plates to recover overnight. The plates were covered by foil to prevent light exposure. About 30 males were placed with 10 anesthetized hermaphrodites (0.1% tricaine and 0.01% tetramisole in M9 buffer) on MYOB plates seeded with a 50-100 μl OP50 bacteria. After 30 minutes of mating, hermaphrodites were then isolated and allowed to rest on food for one hour. The mated hermaphrodites were then mounted for microscopy on 5% agarose pads with the anesthetic. Image acquisition was captured by a Nikon 60 X 1.2 NA water objective with 1 um z-step size. Quantification of sperm distribution in the uterus starts at the vulva and extends up to and includes the spermatheca. The whole uterus was divided into three zones. Zone 1 contains the vulva region, and Zone 3 contains the spermatheca. The number of sperm was manually counted within each zone. The distribution percentage= (the number in each zone) / (the total labeled sperm observed) * 100%. The quantified data contains at least 30 total stained sperm in the entire uterus. At least 3-7 mated hermaphrodites were counted in each mating assay, and experiments were repeated at least 3 times.

### Auxin-inducible treatment in the degron strains

Animals were grown on bacteria-seeded MYOB plates containing auxin. The natural auxin indole-3-acetic acid (IAA) was purchased from Alfa Aesar (#A10556). IAA was dissolved in ethanol as a 400 mM stock solution. Auxin was added to autoclaved MYOB agar when it cooled to about 50-60°C before pouring. MYOB plates containing the final concentration of auxin (1 or 2 mM) were used to test the degron-edited worms. To efficiently degrade the target protein, L1 or L2 hermaphrodites were picked onto auxin plates. Animals were grown on the plates at 20°C for 36-60 hours for the brood size assay. Alternatively, mid-L4 hermaphrodites were incubated on the auxin plate for one generation, and F1 mid-L4 hermaphrodites were picked to a fresh auxin plate for the brood size assay or phenotypic imaging.

### Microscopy

Live imaging was performed on a spinning disk confocal system that uses a Nikon 60 X 1.2 NA water objective, a Photometrics Prime 95B EMCCD camera, and a Yokogawa CSU-X1 confocal scanner unit. Images were acquired and analyzed by Nikon’s NIS imaging software and ImageJ/FIJI Bio-formats plugin (National Institutes of Health) (Linkert et al., 2010; Schindelin et al., 2012). GCaMP3 images were also acquired by a 60×/1.40 NA oil-immersion objective on a Nikon Eclipse 80i microscope equipped with a SPOT RT39M5 sCMOS camera (Diagnostic Instruments, Sterling Heights, MI, USA) with a 0.63x wild field adapter, controlled by SPOT Advanced imaging software (v. 5.0) with Peripheral Devices and Quantitative Imaging modules. Images were acquired at 2448 × 2048 pixels, using the full camera chip, and saved as 8-bit TIFF files. Fluorescence excitation was provided by a Nikon Intensilight C-HGFI 130-W mercury lamp and shuttered with a Lambda 10-B SmartShutter (Sutter Instruments, Novato, CA), also controlled through the SPOT software. Single-channel GCaMP time-lapse movies were acquired using a GFP filter set (470/40× 495lpxr 525/50m) (Chroma Technologies, Bellows Falls VT) at 1 frame per second, with an exposure time of 40-60 ms, gain of 8, and neutral density of 16.

### GCaMP3 imaging acquisition and data processing

For all GCaMP3 imaging data, animals were immobilized on 7.5% agarose pads with 0.05 μm polystyrene beads and imaged using confocal microscopy as described above. Images were acquired every 1 second and saved as 16-bit TIFF files. DIC images were acquired every 3 seconds. Only successful embryo transits (embryos that were expelled through the sp-ut valve) were analyzed for this GCaMP3 study. The GCaMP3 metrics, including rising time and fraction over half max data, as well as the GCaMP3 intensity heat map were processed by the custom Fiji and Matlab coded platform (Bouffard et al., 2019). GCaMP3 kymograms were generated by custom Fiji code using the commands Image>Stacks>Reslice followed by Image>Stacks>Z Project (Average Intensity) (Bouffard et al., 2019). Only the very first three ovulations were imaged for each animal. Detailed Processing and analysis of the GCaMP time series was performed exactly as described in (Bouffard et al., 2019).

### Statistics

Statistical significance was determined by p value from an unpaired 2 tailed t-test. P-values: ns = not significant; * = <0.05; ** = <0.01, *** = <0.001; **** = <0.0001. Both the Shapiro-Wilk and Kolmogorov-Smirnov Normality test indicated that all data follow normal distributions.

## Acknowledgments

We thank the *Caenorhabditis* Genetics Center, which is funded by National Institutes of Health Office of Research Infrastructure Programs (P40OD010440), for providing strains for this study. We thank Dr. Orna Cohen-Fix for generously sharing the SP-12::GFP strain, and Dr. Harold Smith for sharing the BA17 *fem-1(hc17ts)* strain. We are grateful to the members of the Golden laboratory, Dr. Peter Kropp, Dr. Tao Cai, Rosie Bauer, Isabella Zafra, and Carina Graham for productive discussions and preparing reagents. We thank our summer intern Kyle Wilson for manuscript editing. We especially thank Dr. Harold Smith, Dr. Orna Cohen-Fix, Dr. Kevin O’ Connell, Dr. Katherine McJunkin and Dan Konzman for critical inputs on the project and feedback on the manuscript. We thank all members of the Baltimore Worm Club for providing feedback and suggestions to our investigations.

## Funding

This work was supported, in part, by the Intramural Research Program of the National Institutes of Health, National Institute of Diabetes and Digestive and Kidney Diseases (X.B. and A.G.), National Institutes of Health, National Institute of General Medical Sciences (GM110268; J.B., A.L. and E.J.C.), and National Institutes of Health grants NIH R24 0D023041 and NIH R01 NS113119 (K.B. and P.W.S).

## Figure Legends

**Supplemental Figure 1. PEZO-1 is expressed in multiple tissues throughout development**

(A) There are 14 mRNA isoforms encoded by *pezo-1*. Isoforms i-l encode the six short forms of *pezo-1* (red asterisks). The 5’-3’ orientation is right to left. (B-G) Both PEZO-1::mScarlet (magenta) and GFP::PEZO-1 (green) express in a variety of cell types, including pharyngeal neurons (B, white arrows), pharyngeal-intestinal valve (C), male tail (D), vulva (E), intestinal cells (F) and seam cells (G). Scale bars are indicated in each panel. Illustration in panel A was taken from WormBase (https://wormbase.org).

**Supplemental Figure 2. Verification of CRISPR/Cas9 generated deletions in *pezo-1* knockout animals**

(A) Representative PCR gel from genotyping single animals for *pezo-1 C-Δ k*nockout candidates. A positive homozygous knockout line is labeled with a red asterisk. Three primers (two that flank the deletion and one internal) were used to test the homozygosity of candidate *pezo-1* deletion animals. Amplicon size of a homozygous deletion with both flanking primers is 450 bp (labeled −/−). In wild type, an 879bp PCR product was able to be amplified by one flanking primer and the internal primer (labelled +/+). Heterozygous animals contain both of the PCR products (labeled +/−). (B) Schematic of the 14 mRNA isoforms and the position of the three deletion alleles used in this study and which isoforms they should affect. The STOP-IN line is also shown as an insertion in the beginning of exon 27. The 5’-3’ orientation is right to left. (C) Two other alleles generated for this study also had reduced brood sizes; both a stop-in mutant *pezo-1 (sy1199)* and a small deletion allele *pezo-1 (sy1398)* in isoforms I and J. (D) The reduction in brood size of *pezo-1* deletion animals was enhanced when the animals were grown at 25°C. (E) Quantification of the percentage of uteri with ooplasmic masses in *pezo-1 (sy1199)* and *pezo-1 (sy1398)* mutants. P-values: *** = 0.0003 (C); ** = 0.0021 (D); *** = 0.0002 (D); **** <0.0001 (*t*-test). Illustration in panel B was taken from WormBase (https://wormbase.org).

**Supplemental Figure 3. Normal calcium signaling was observed in the spermathecal cells in *pezo-1* mutants**

(A, B) GCaMP3 time series of normalized average pixel intensity from a single oocyte transit recording over the same spatial frame and time. (C) Heat map of GCaMP3 normalized average pixel intensity (F/F0) versus time series during ovulation from seven oocyte transit recordings in both wildtype and *pezo-1 C-Δ* mutants. Color bars represents the gradient of the normalized average pixel intensity (F/F0). (D, E) Representative kymograms of GCaMP3 in both wildtype and *pezo-1 C-Δ* mutants. Color bars represents the gradient of the fluorescence intensity.

**Supplemental Figure 4. Somatic-tissue specific degradation of PEZO-1 causes severe ovulation defects**

(A-H) Abnormal ovulations were observed in the somatic tissue specific PEZO-1::Degron animals. Shown are two different ovulation events. (A, E) Ovulation initiated by oocyte (yellow dotted circle) entry into the spermatheca. Spermathecal distal valve (red arrows) was defective (B, C, E-H) and either pinched off the oocyte when it attempted to enter the spermatheca (B-D) or failed to open and block/delayed the entry of the oocyte into the spermatheca (yellow asterisks) (E-H). Timing of each step is labeled in each panel in minutes and seconds. (I) Quantification of the oocyte ovulation rate and ovulation defects in the *Peft-3::tir-1; pezo-1::Degron* animals with or without 2 mM auxin. Scale bars are indicated in each panel.

**Supplemental Figure 5. *pezo-1* mutants genetically interact with the gap junctional proteins**

(A, B) Quantification of brood size and embryonic viability of wildtype and *pezo-1* mutants after RNAi feeding of *inx-14* and *inx-22*. P-values: * = 0.018 (A); ** = 0.0019 (A); *** =0.0006 (B); **** <0.0001 (*t*-test).

**Supplemental Figure 6. Excessive yolk was observed in *pezo-1* mutants**

(A-C’) A vitellogenin::tdimer2 fluorescent protein (YP170::tdimer2) transgene was used to monitor yolk lipoprotein localization. (B, C, B’, C’) Excess yolk was observed in the pseudocoelom of *pezo-1* mutants (white dotted regions), suggesting that yolk endocytosis is partially defective.

**Video S1. PEZO-1 expression pattern during ovulation**

Ovulation imaged in the genome-edited animals expressing GFP::PEZO-1 (green). Yellow arrow in right panel indicates GFP::PEZO-1 expression on the spermathecal valves. White arrows in right panel indicate GFP::PEZO-1 expression on the bag cells. After fertilization, GFP::PEZO-1 labeled sperm crawled back to the spermatheca. Left panel shows the merged channel of DIC (grey) with GFP (green). Right panel indicates the GFP (green) channel only. Images are single z planes taken every 2 seconds. Timing is indicated in lower right. Playback rate is 15 frames/second. Scale bar is indicated in left panel.

**Video S2. Crushed oocyte phenotype frequently occurs in the *pezo-1 C-Δ* mutant**

Time-lapse video recording showing a wildtype oocyte (top panel) entering into the spermatheca and completing fertilization in 5 minutes. The constricted spermatheca smoothly expels the oocyte into the uterus. White arrows in top panel indicate opening spermathecal valve. In the bottom panel, the *pezo-1 C-Δ* oocyte successfully enters the spermatheca, but the oocyte is crushed by the sp-ut valve and the ooplasmic debris is observed in the uterus. Yellow arrows in bottom panel indicate the spermathecal valve. Images are single z planes taken every 2 seconds. Timing is indicated in lower right. Playback rate is 15 frames/second. Scale bars are indicated in each panel.

**Video S3. The sp-ut valve fails to open during spermathecal contraction**

Time-lapse recordings on left are of DIC and GFP. Recordings on right are only of GFP. Oocyte entry occurs from the left at the 15 second mark. The spermatheca was labelled by the apical junctional marker DLG-1::GFP. In the wild type (top panels), the sp-ut valve (white arrow) opened immediately to allow the oocyte to be expelled into the uterus (on right). However, in *pezo-1 C-Δ* (bottom panels), the DLG-1::GFP labelled sp-ut valve (white arrow) never fully opened, the oocyte was crushed as it was expelled, and ooplasmic debris was pushed out into the uterus. Images are single z planes taken every 3 seconds. Timing is indicated in lower right. Playback rate is 15 frames/second. Scale bars are indicated in each DIC panel.

**Video S4. Spermatheca dilation is defective in *pezo-1* mutants**

Time-lapse video recording (DIC). Oocyte entry occurs from the left at the 35 second mark. The distal valve was not able to completely close and the oocyte was pinched. One portion of the broken oocyte was left in the spermatheca, the other portion remains in the oviduct (white arrows, left panel). Images are single z planes taken every 2 seconds. Timing is indicated in lower left. Playback rate is 15 frames/second. Scale bar is indicated in lower right.

**Video S5. Sheath cell contraction is defective in *pezo-1* mutants**

Time-lapse video recording (DIC). Oocyte that fails to enter the spermatheca after a few attempts. Sheath cells failed to contract and push the oocyte into the spermatheca (on the right) and oocyte moves left, back into the oviduct. Images are single z planes taken every 2 seconds. Timing is indicated in lower right. Playback rate is 15 frames/second. Scale bar is indicated in lower left.

**Video S6. mScarlet::PEZO-1 colocalizes with spermathecal-specific GCaMP3**

Example of the colocalization of mScarlet::PEZO-1 (magenta) with the Pfln-1::GCaMP3 transgene (green) in the spermathecal cells in a wildtype animal. Top left recording shows the merged channel of DIC (grey), mScarlet::PEZO-1 (magenta) and the *Pfln-1::GCaMP3* transgene (green). Top right panel lacks the DIC channel. Bottom left recording shows just the mScarlet::PEZO-1 expression pattern during ovulation. Bottom right indicates that Pfln-1::GCaMP3 only displays the changes of GCaMP3 intensity, which is indicative of calcium influx. Images were acquired in a single z plane every 2 seconds. Timing is indicated in lower right panel. Playback rate is 30 frames/second. Scale bars are indicated in each panel.

**Video S7. Normal GCaMP3 influx was observed in *pezo-1* mutants**

Examples of GCaMP3 recordings of embryo transits in wildtype (left panels) and *pezo-1 C-Δ* (right panels) animals. Recordings were temporally aligned to the start of oocyte entry at 50 seconds. GCaMP3 normalized average pixel intensity (F/F0, top, Y-axis) versus GCaMP3 time (top, X-axis) generated from GCaMP3 recordings with highlighted metrics on the top of the tracings. Dwell time is a tissue function metric that represents the duration from the closing of the distal valve to the opening of the sp-ut valve, rising time is a calcium signaling metric measuring the time from the opening of the distal valve to the first time point where the time series reaches half maximum of GCaMP3 intensity, and fraction over half max is a calcium signaling metric, which measures the duration of the dwell time over the GCaMP3 half-maximal value divided by the total dwell time. Images were acquired in a single z plane every 1 second. Timing is indicated in top left of each bottom panel. Playback rate is 30 frames/second. Scale bars are indicated in each panel.

## Bibliography

Albuisson, J., Murthy, S.E., Bandell, M., Coste, B., Louis-Dit-Picard, H., Mathur, J., Feneant-Thibault, M., Tertian, G., de Jaureguiberry, J.P., Syfuss, P.Y., Cahalan, S., Garcon, L., Toutain, F., Simon Rohrlich, P., Delaunay, J., Picard, V., Jeunemaitre, X., Patapoutian, A., 2013. Dehydrated hereditary stomatocytosis linked to gain-of-function mutations in mechanically activated PIEZO1 ion channels. Nat Commun 4, 1884.

Alper, S.L., 2017. Genetic Diseases of PIEZO1 and PIEZO2 Dysfunction. Curr Top Membr 79, 97–134.

Andolfo, I., Alper, S.L., De Franceschi, L., Auriemma, C., Russo, R., De Falco, L., Vallefuoco, F., Esposito, M.R., Vandorpe, D.H., Shmukler, B.E., Narayan, R., Montanaro, D., D’Armiento, M., Vetro, A., Limongelli, I., Zuffardi, O., Glader, B.E., Schrier, S.L., Brugnara, C., Stewart, G.W., Delaunay, J., Iolascon, A., 2013. Multiple clinical forms of dehydrated hereditary stomatocytosis arise from mutations in PIEZO1. Blood 121, 3925–3935, S3921-3912.

Bae, C., Gnanasambandam, R., Nicolai, C., Sachs, F., Gottlieb, P.A., 2013. Xerocytosis is caused by mutations that alter the kinetics of the mechanosensitive channel PIEZO1. P Natl Acad Sci USA 110, E1162–E1168.

Bouffard, J., Cecchetelli, A.D., Clifford, C., Sethi, K., Zaidel-Bar, R., Cram, E.J., 2019. The RhoGAP SPV-1 regulates calcium signaling to control the contractility of the Caenorhabditis elegans spermatheca during embryo transits. Mol Biol Cell 30, 907–922.

Bui, Y.K., Sternberg, P.W., 2002. Caenorhabditis elegans inositol 5-phosphatase homolog negatively regulates inositol 1,4,5-triphosphate signaling in ovulation. Mol Biol Cell 13, 1641–1651.

Clandinin, T.R., DeModena, J.A., Sternberg, P.W., 1998. Inositol trisphosphate mediates a RAS-independent response to LET-23 receptor tyrosine kinase activation in C. elegans. Cell 92, 523–533.

Coste, B., Houge, G., Murray, M.F., Stitziel, N., Bandell, M., Giovanni, M.A., Philippakis, A., Hoischen, A., Riemer, G., Steen, U., Steen, V.M., Mathur, J., Cox, J., Lebo, M., Rehm, H., Weiss, S.T., Wood, J.N., Maas, R.L., Sunyaev, S.R., Patapoutian, A., 2013. Gain-of-function mutations in the mechanically activated ion channel PIEZO2 cause a subtype of Distal Arthrogryposis. Proc Natl Acad Sci U S A 110, 4667–4672.

Coste, B., Mathur, J., Schmidt, M., Earley, T.J., Ranade, S., Petrus, M.J., Dubin, A.E., Patapoutian, A., 2010. Piezo1 and Piezo2 are essential components of distinct mechanically activated cation channels. Science 330, 55–60.

Coste, B., Xiao, B., Santos, J.S., Syeda, R., Grandl, J., Spencer, K.S., Kim, S.E., Schmidt, M., Mathur, J., Dubin, A.E., Montal, M., Patapoutian, A., 2012. Piezo proteins are pore-forming subunits of mechanically activated channels. Nature 483, 176–181.

Cram, E.J., 2014. Mechanotransduction in C. elegans morphogenesis and tissue function. Prog Mol Biol Transl Sci 126, 281–316.

Cram, E.J., 2015. Mechanotransduction: feeling the squeeze in the C. elegans reproductive system. Curr Biol 25, R74–R75.

Del Marmol, J.I., Touhara, K.K., Croft, G., MacKinnon, R., 2018. Piezo1 forms a slowly-inactivating mechanosensory channel in mouse embryonic stem cells. Elife 7.

Doniach, T., Hodgkin, J., 1984. A Sex-Determining Gene, Fem-1, Required for Both Male and Hermaphrodite Development in Caenorhabditis-Elegans. Developmental Biology 106, 223–235.

Edmonds, J.W., McKinney, S.L., Prasain, J.K., Miller, M.A., 2011. The gap junctional protein INX-14 functions in oocyte precursors to promote C. elegans sperm guidance. Dev Biol 359, 47–58.

Fraser, A.G., Kamath, R.S., Zipperlen, P., Martinez-Campos, M., Sohrmann, M., Ahringer, J., 2000. Functional genomic analysis of C. elegans chromosome I by systematic RNA interference. Nature 408, 325–330.

Friedland, A.E., Tzur, Y.B., Esvelt, K.M., Colaiacovo, M.P., Church, G.M., Calarco, J.A., 2013. Heritable genome editing in C. elegans via a CRISPR-Cas9 system. Nat Methods 10, 741–743.

Gnanasambandam, R., Bae, C., Gottlieb, P.A., Sachs, F., 2015. Ionic Selectivity and Permeation Properties of Human PIEZO1 Channels. PLoS One 10, e0125503.

Govindan, J.A., Nadarajan, S., Kim, S., Starich, T.A., Greenstein, D., 2009. Somatic cAMP signaling regulates MSP-dependent oocyte growth and meiotic maturation in C. elegans. Development 136, 2211–2221.

Greenstein, D., 2005. Control of oocyte meiotic maturation and fertilization. WormBook, 1–12.

Han, S.M., Cottee, P.A., Miller, M.A., 2010. Sperm and oocyte communication mechanisms controlling C. elegans fertility. Dev Dyn 239, 1265–1281.

Harris, T.W., Arnaboldi, V., Cain, S., Chan, J., Chen, W.J., Cho, J., Davis, P., Gao, S., Grove, C.A., Kishore, R., Lee, R.Y.N., Muller, H.M., Nakamura, C., Nuin, P., Paulini, M., Raciti, D., Rodgers, F.H., Russell, M., Schindelman, G., Auken, K.V., Wang, Q., Williams, G., Wright, A.J., Yook, K., Howe, K.L., Schedl, T., Stein, L., Sternberg, P.W., 2019. WormBase: a modern Model Organism Information Resource. Nucleic Acids Res.

He, L., Si, G., Huang, J., Samuel, A.D.T., Perrimon, N., 2018. Mechanical regulation of stem-cell differentiation by the stretch-activated Piezo channel. Nature 555, 103–106.

Hoang, H.D., Prasain, J.K., Dorand, D., Miller, M.A., 2013. A heterogeneous mixture of F-series prostaglandins promotes sperm guidance in the Caenorhabditis elegans reproductive tract. PLoS Genet 9, e1003271.

Kelley, C.A., Cram, E.J., 2019. Regulation of Actin Dynamics in the C. elegans Somatic Gonad. J Dev Biol 7.

Kim, S.E., Coste, B., Chadha, A., Cook, B., Patapoutian, A., 2012. The role of Drosophila Piezo in mechanical nociception. Nature 483, 209–212.

Kimble, J., Hirsh, D., 1979. The postembryonic cell lineages of the hermaphrodite and male gonads in Caenorhabditis elegans. Dev Biol 70, 396–417.

Kovacevic, I., Orozco, J.M., Cram, E.J., 2013. Filamin and phospholipase C-epsilon are required for calcium signaling in the Caenorhabditis elegans spermatheca. PLoS Genet 9, e1003510.

Kubagawa, H.M., Watts, J.L., Corrigan, C., Edmonds, J.W., Sztul, E., Browse, J., Miller, M.A., 2006. Oocyte signals derived from polyunsaturated fatty acids control sperm recruitment in vivo. Nat Cell Biol 8, 1143–1148.

Kuwabara, P.E., 2003. The multifaceted C-elegans major sperm protein: an ephrin signaling antagonist in oocyte maturation. Gene Dev 17, 155–161.

Li, J., Hou, B., Tumova, S., Muraki, K., Bruns, A., Ludlow, M.J., Sedo, A., Hyman, A.J., McKeown, L., Young, R.S., Yuldasheva, N.Y., Majeed, Y., Wilson, L.A., Rode, B., Bailey, M.A., Kim, H.R., Fu, Z., Carter, D.A., Bilton, J., Imrie, H., Ajuh, P., Dear, T.N., Cubbon, R.M., Kearney, M.T., Prasad, R.K., Evans, P.C., Ainscough, J.F., Beech, D.J., 2014. Piezo1 integration of vascular architecture with physiological force. Nature 515, 279–282.

Li, J., Hou, B., Tumova, S., Muraki, K., Bruns, A., Ludlow, M.J., Sedo, A., Hyman, A.J., McKeown, L., Young, R.S., Yuldasheva, N.Y., Majeed, Y., Wilson, L.A., Rode, B., Bailey, M.A., Kim, H.R., Fu, Z.J., Carter, D.A.L., Bilton, J., Imrie, H., Ajuh, P., Dear, T.N., Cubbon, R.M., Kearney, M.T., Prasad, K.R., Evans, P.C., Ainscough, J.F.X., Beech, D.J., 2015. Piezo1 Integration of Vascular Architecture with Physiological Force. Faseb J 29.

Li, S., You, Y., Gao, J., Mao, B., Cao, Y., Zhao, X., Zhang, X., 2018. Novel mutations in TPM2 and PIEZO2 are responsible for distal arthrogryposis (DA) 2B and mild DA in two Chinese families. BMC Med Genet 19, 179.

Linkert, M., Rueden, C.T., Allan, C., Burel, J.M., Moore, W., Patterson, A., Loranger, B., Moore, J., Neves, C., Macdonald, D., Tarkowska, A., Sticco, C., Hill, E., Rossner, M., Eliceiri, K.W., Swedlow, J.R., 2010. Metadata matters: access to image data in the real world. J Cell Biol 189, 777–782.

Lorin-Nebel, C., Xing, J., Yan, X.H., Strange, K., 2007. CRAC channel activity in C-elegans is mediated by Orai1 and STIM1 homologues and is essential for ovulation and fertility. J Physiol-London 580, 67–85.

Lukacs, V., Mathur, J., Mao, R., Bayrak-Toydemir, P., Procter, M., Cahalan, S.M., Kim, H.J., Bandell, M., Longo, N., Day, R.W., Stevenson, D.A., Patapoutian, A., Krock, B.L., 2015. Impaired PIEZO1 function in patients with a novel autosomal recessive congenital lymphatic dysplasia. Nat Commun 6, 8329.

McCarter, J., Bartlett, B., Dang, T., Schedl, T., 1999. On the control of oocyte meiotic maturation and ovulation in Caenorhabditis elegans. Dev Biol 205, 111–128.

McHugh, B.J., Buttery, R., Lad, Y., Banks, S., Haslett, C., Sethi, T., 2010. Integrin activation by Fam38A uses a novel mechanism of R-Ras targeting to the endoplasmic reticulum. J Cell Sci 123, 51–61.

McKnight, K., Hoang, H.D., Prasain, J.K., Brown, N., Vibbert, J., Hollister, K.A., Moore, R., Ragains, J.R., Reese, J., Miller, M.A., 2014. Neurosensory perception of environmental cues modulates sperm motility critical for fertilization. Science 344, 754–757.

McMillin, M.J., Beck, A.E., Chong, J.X., Shively, K.M., Buckingham, K.J., Gildersleeve, H.I., Aracena, M.I., Aylsworth, A.S., Bitoun, P., Carey, J.C., Clericuzio, C.L., Crow, Y.J., Curry, C.J., Devriendt, K., Everman, D.B., Fryer, A., Gibson, K., Giovannucci Uzielli, M.L., Graham, J.M., Jr., Hall, J.G., Hecht, J.T., Heidenreich, R.A., Hurst, J.A., Irani, S., Krapels, I.P., Leroy, J.G., Mowat, D., Plant, G.T., Robertson, S.P., Schorry, E.K., Scott, R.H., Seaver, L.H., Sherr, E., Splitt, M., Stewart, H., Stumpel, C., Temel, S.G., Weaver, D.D., Whiteford, M., Williams, M.S., Tabor, H.K., Smith, J.D., Shendure, J., Nickerson, D.A., University of Washington Center for Mendelian, G., Bamshad, M.J., 2014. Mutations in PIEZO2 cause Gordon syndrome, Marden-Walker syndrome, and distal arthrogryposis type 5. Am J Hum Genet 94, 734–744.

Miller, M.A., 2001. A sperm cytoskeletal protein that signals oocyte meiotic maturation and ovulation (vol 292, pg 2144, 2001). Science 292, 639–639.

Miller, M.A., Ruest, P.J., Kosinski, M., Hanks, S.K., Greenstein, D., 2003. An Eph receptor sperm-sensing control mechanism for oocyte meiotic maturation in Caenorhabditis elegans. Genes Dev 17, 187–200.

Murthy, S.E., Dubin, A.E., Patapoutian, A., 2017. Piezos thrive under pressure: mechanically activated ion channels in health and disease. Nat Rev Mol Cell Biol 18, 771–783.

Nonomura, K., Lukacs, V., Sweet, D.T., Goddard, L.M., Kanie, A., Whitwam, T., Ranade, S.S., Fujimori, T., Kahn, M.L., Patapoutian, A., 2018. Mechanically activated ion channel PIEZO1 is required for lymphatic valve formation. P Natl Acad Sci USA 115, 12817–12822.

Nonomura, K., Woo, S.H., Chang, R.B., Gillich, A., Qiu, Z., Francisco, A.G., Ranade, S.S., Liberles, S.D., Patapoutian, A., 2017. Piezo2 senses airway stretch and mediates lung inflation-induced apnoea. Nature 541, 176–181.

Paix, A., Folkmann, A., Rasoloson, D., Seydoux, G., 2015. High Efficiency, Homology-Directed Genome Editing in Caenorhabditis elegans Using CRISPR-Cas9 Ribonucleoprotein Complexes. Genetics 201, 47-+.

Periasamy, M., Huke, S., 2001. SERCA pump level is a critical determinant of Ca2+ homeostasis and cardiac contractility. J Mol Cell Cardiol 33, 1053–1063.

Poole, K., Herget, R., Lapatsina, L., Ngo, H.D., Lewin, G.R., 2014. Tuning Piezo ion channels to detect molecular-scale movements relevant for fine touch. Nat Commun 5, 3520.

Ranade, S.S., Qiu, Z.Z., Woo, S.H., Hur, S.S., Murthy, S.E., Cahalan, S.M., Xu, J., Mathur, J., Bandell, M., Coste, B., Li, Y.S.J., Chien, S., Patapoutian, A., 2014. Piezo1, a mechanically activated ion channel, is required for vascular development in mice. P Natl Acad Sci USA 111, 10347–10352.

Rual, J.F., Ceron, J., Koreth, J., Hao, T., Nicot, A.S., Hirozane-Kishikawa, T., Vandenhaute, J., Orkin, S.H., Hill, D.E., van den Heuvel, S., Vidal, M., 2004. Toward improving Caenorhabditis elegans phenome mapping with an ORFeome-based RNAi library. Genome Res 14, 2162–2168.

Schindelin, J., Arganda-Carreras, I., Frise, E., Kaynig, V., Longair, M., Pietzsch, T., Preibisch, S., Rueden, C., Saalfeld, S., Schmid, B., Tinevez, J.Y., White, D.J., Hartenstein, V., Eliceiri, K., Tomancak, P., Cardona, A., 2012. Fiji: an open-source platform for biological-image analysis. Nat Methods 9, 676–682.

Singson, A., Mercer, K.B., L’Hernault, S.W., 1998. The C-elegans spe-9 gene encodes a sperm transmembrane protein that contains EGF-like repeats and is required for fertilization. Cell 93, 71–79.

Syeda, R., Xu, J., Dubin, A.E., Coste, B., Mathur, J., Huynh, T., Matzen, J., Lao, J., Tully, D.C., Engels, I.H., Petrassi, H.M., Schumacher, A.M., Montal, M., Bandell, M., Patapoutian, A., 2015. Chemical activation of the mechanotransduction channel Piezo1. Elife 4.

Vicencio, J., Martinez-Fernandez, C., Serrat, X., Ceron, J., 2019. Efficient Generation of Endogenous Fluorescent Reporters by Nested CRISPR in Caenorhabditis elegans. Genetics 211, 1143–1154.

Voglis, G., Tavernarakis, N., 2005. Mechanotransduction in the Nematode Caenorhabditis elegans, in: Kamkin, A., Kiseleva, I. (Eds.), Mechanosensitivity in Cells and Tissues, Moscow.

Whitten, S.J., Miller, M.A., 2007. The role of gap junctions in Caenorhabditis elegans oocyte maturation and fertilization. Developmental Biology 301, 432–446.

Woo, S.H., Lukacs, V., de Nooij, J.C., Zaytseva, D., Criddle, C.R., Francisco, A., Jessell, T.M., Wilkinson, K.A., Patapoutian, A., 2015. Piezo2 is the principal mechanotransduction channel for proprioception. Nat Neurosci 18, 1756–1762.

Woo, S.H., Ranade, S., Weyer, A.D., Dubin, A.E., Baba, Y., Qiu, Z.Z., Petrus, M., Miyamoto, T., Reddy, K., Lumpkin, E.A., Stucky, C.L., Patapoutian, A., 2014. Piezo2 is required for Merkel-cell mechanotransduction. Nature 509, 622–626.

Wu, J., Lewis, A.H., Grandl, J., 2017. Touch, Tension, and Transduction - The Function and Regulation of Piezo Ion Channels. Trends Biochem Sci 42, 57–71.

Yan, X., Xing, J., Lorin-Nebel, C., Estevez, A.Y., Nehrke, K., Lamitina, T., Strange, K., 2006. Function of a STIM1 homologue in C. elegans: evidence that store-operated Ca2+ entry is not essential for oscillatory Ca2+ signaling and ER Ca2+ homeostasis. J Gen Physiol 128, 443–459.

Yang, Y., Han, S.M., Miller, M.A., 2010. MSP hormonal control of the oocyte MAP kinase cascade and reactive oxygen species signaling. Dev Biol 342, 96–107.

Zarychanski, R., Schulz, V.P., Houston, B.L., Maksimova, Y., Houston, D.S., Smith, B., Rinehart, J., Gallagher, P.G., 2012. Mutations in the mechanotransduction protein PIEZO1 are associated with hereditary xerocytosis. Blood 120, 1908–1915.

Zhang, L., Ward, J.D., Cheng, Z., Dernburg, A.F., 2015. The auxin-inducible degradation (AID) system enables versatile conditional protein depletion in C. elegans. Development 142, 4374–4384.

Zhang, M., Wang, Y., Geng, J., Zhou, S., Xiao, B., 2019. Mechanically Activated Piezo Channels Mediate Touch and Suppress Acute Mechanical Pain Response in Mice. Cell Rep 26, 1419–1431 e1414.

Zhang, T., Chi, S., Jiang, F., Zhao, Q., Xiao, B., 2017. A protein interaction mechanism for suppressing the mechanosensitive Piezo channels. Nat Commun 8, 1797.

Zwaal, R.R., Van Baelen, K., Groenen, J.T.M., van Geel, A., Rottiers, V., Kaletta, T., Dode, L., Raeymaekers, L., Wuytack, F., Bogaert, T., 2001. The sarco-endoplasmic reticulum Ca2+ ATPase is required for development and muscle function in Caenorhabditis elegans. J Biol Chem 276, 43557–43563.

