## Supplemental Figures for "*Caenorhabditis elegans* PIEZO Channel Coordinates Multiple Reproductive Tissues to Govern Ovulation"

### Supplemental Figure 1

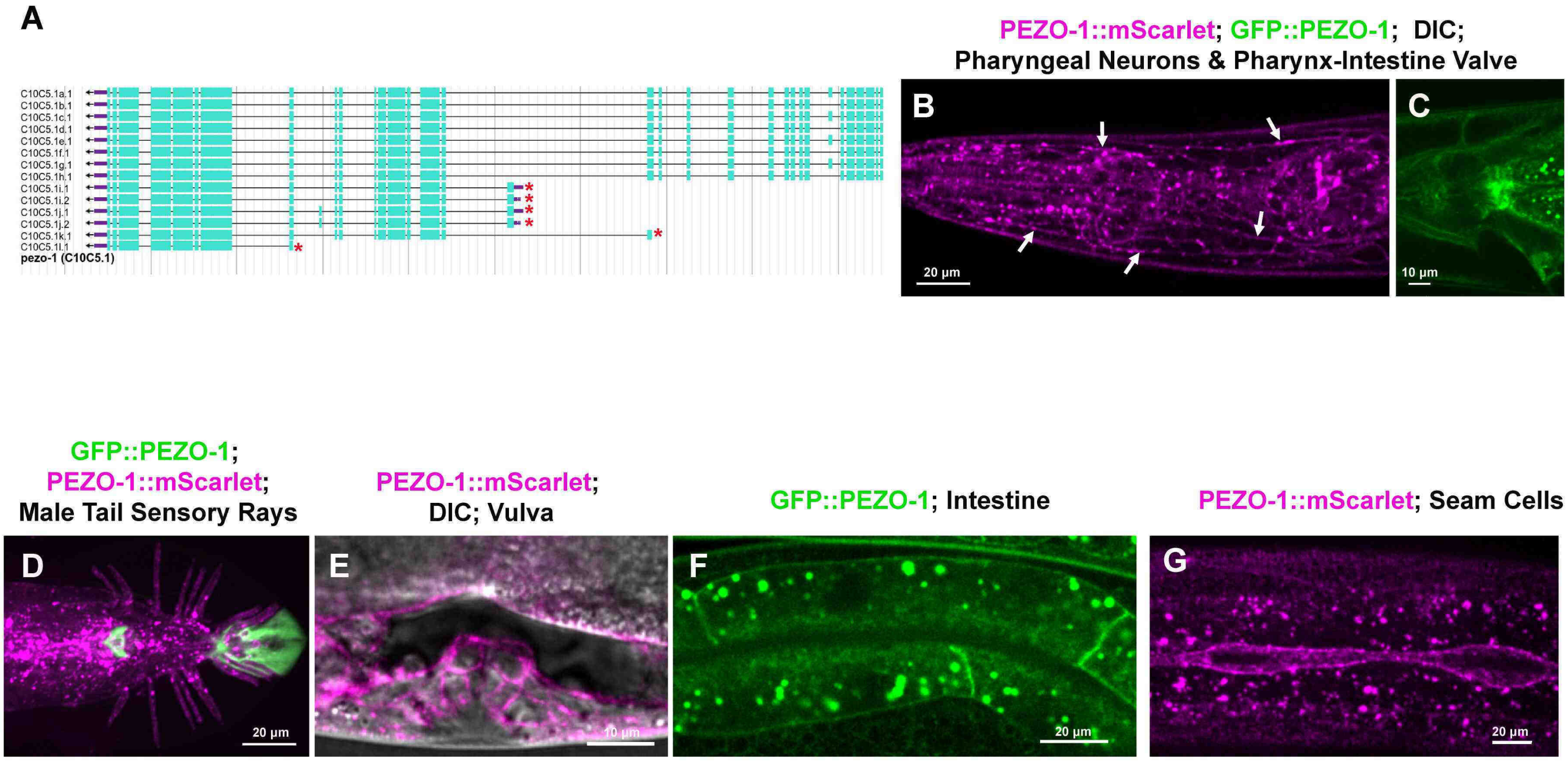

### Supplemental Figure 2

A

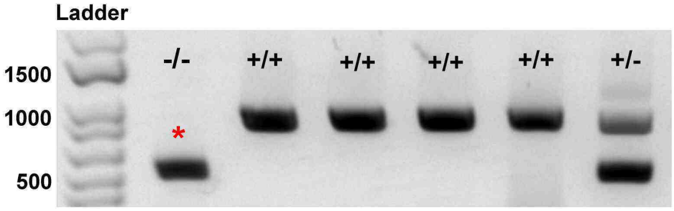


B

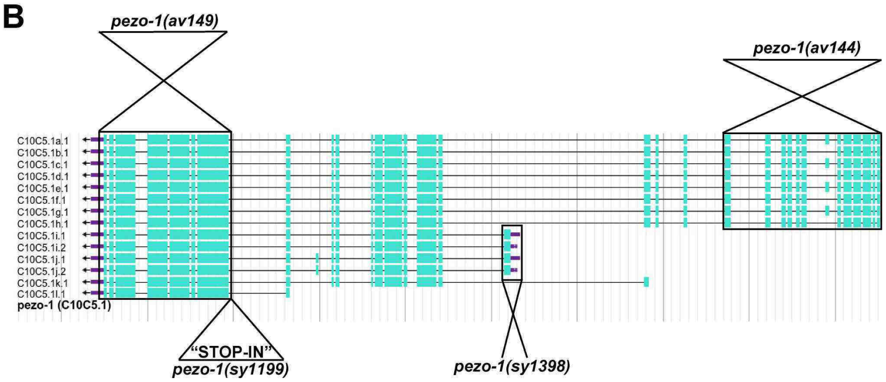


C

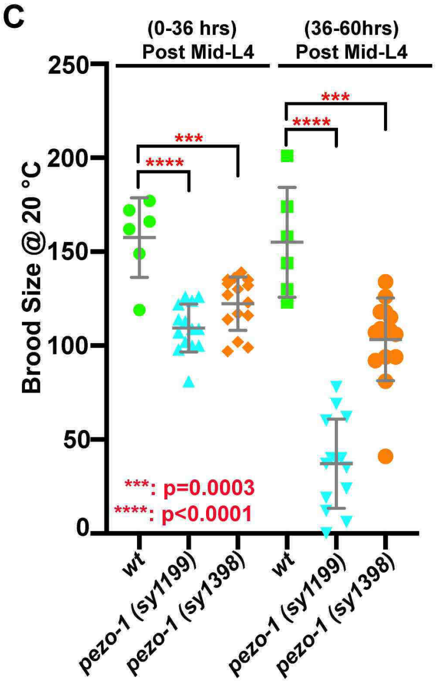


D

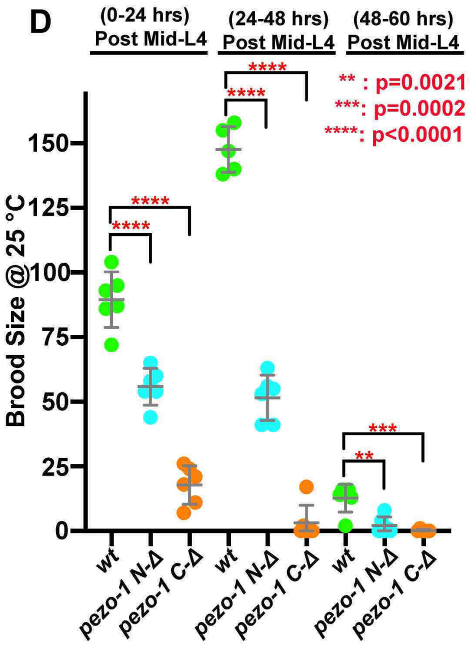


E

| Genotype | # of hrs Post Mid-L4 | Uteri with Ooplasmic Mass |
|---|---|---|
| *pezo-1 (sy1199)* | 36 hours | 50.0% (7/14) |
| | 60 hours | 100% (14/14) |
| *pezo-1 (sy1398)* | 36 hours | 6.7% (1/15) |
| | 60 hours | 26.7% (4/15) |

#### Supplemental Figure 3

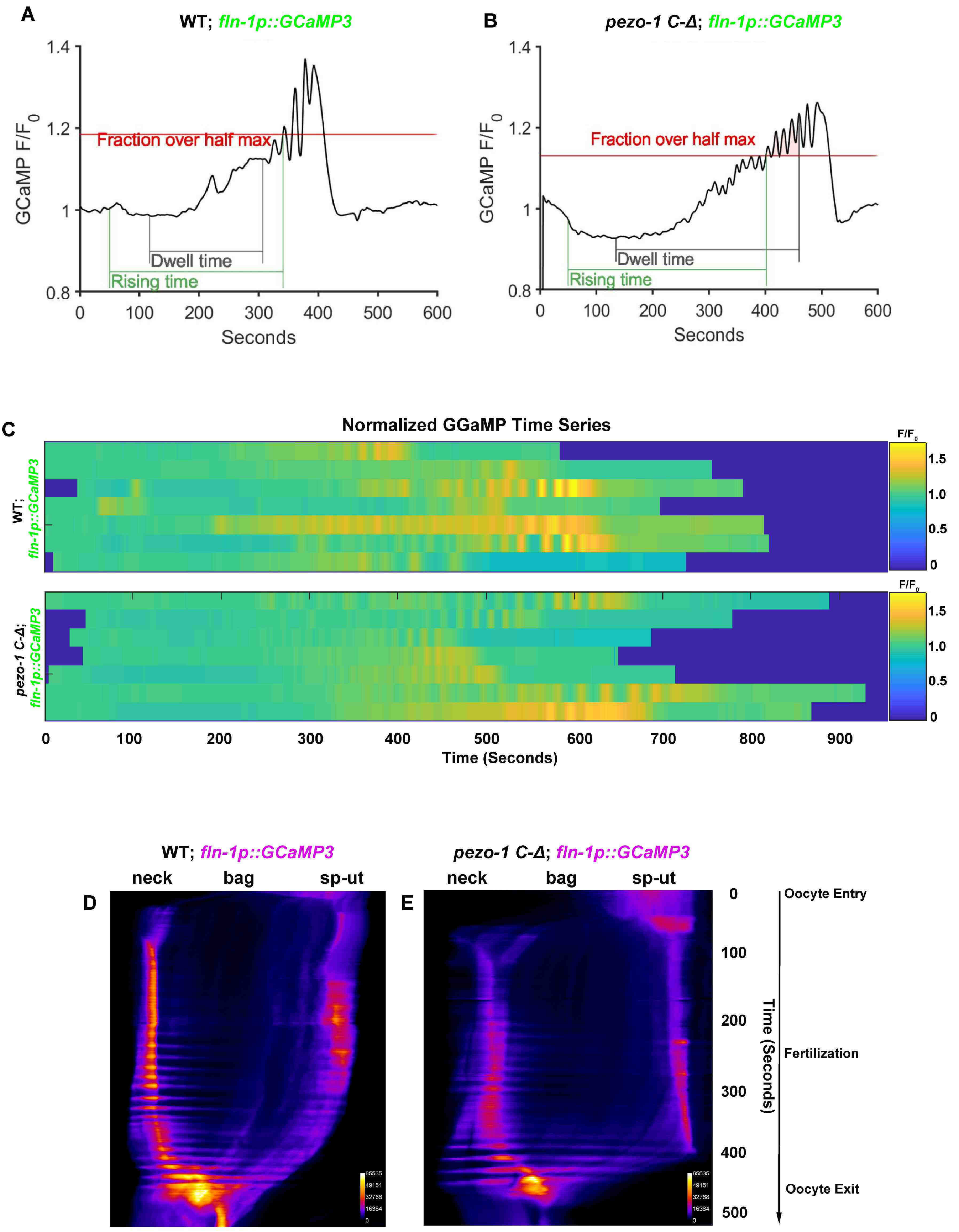

### Supplemental Figure 4

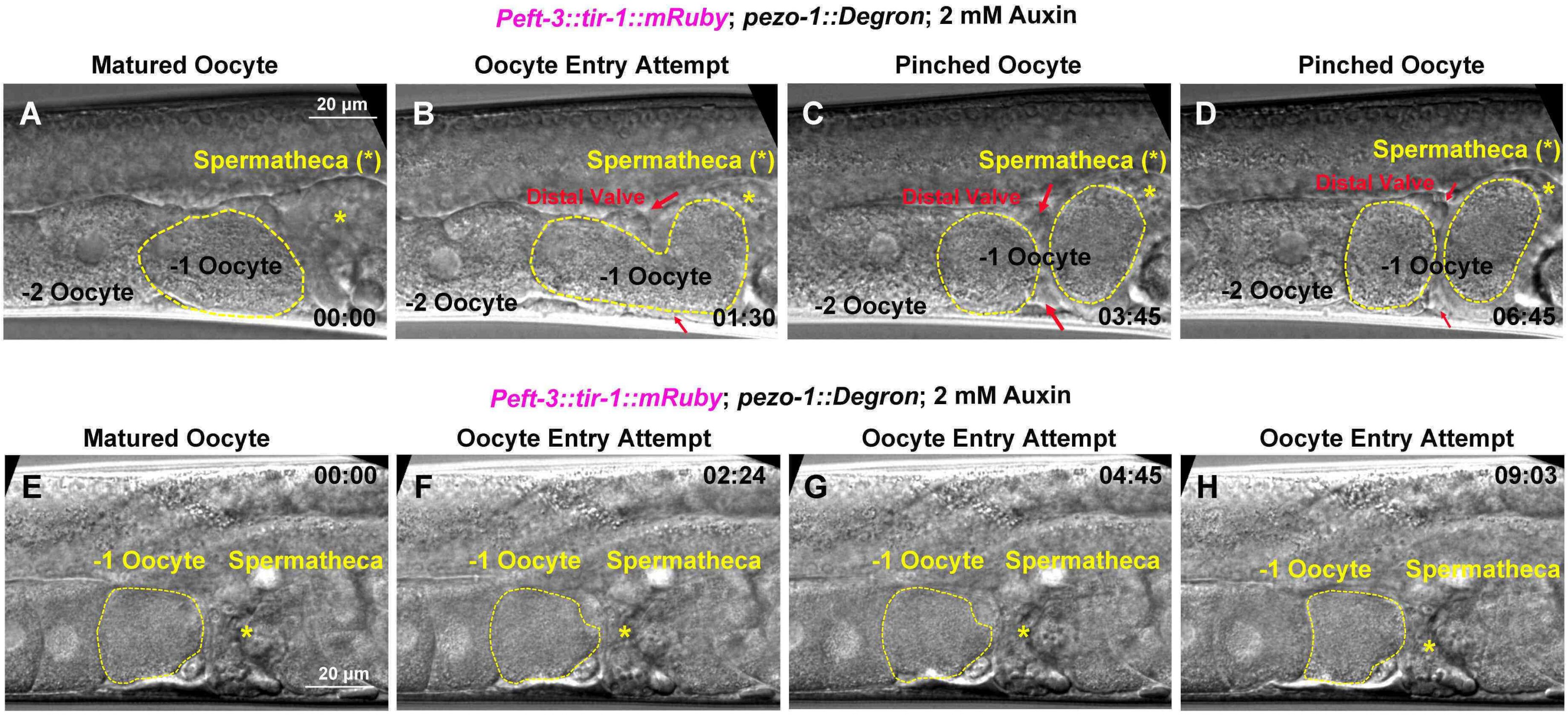


**I**

| Genotype | Treatment | # Ovulations per hour | # Tested Gonad | # Pinched Oocyte | # Delayed Oocyte Entry |
|---|---|---|---|---|---|
| *Peft-3::tir-1::mRuby; pezo-1::Degron* | Control | 1.71 ± 0.49 | 12 | 0% (0/13) | 0% (0/13) |
| *Peft-3::tir-1::mRuby; pezo-1::Degron* | 2 mM Auxin | 1.14 ± 0.65 | 21 | 33.3% (9/27) | 11.1% (3/27) |

#### Supplemental Figure 5

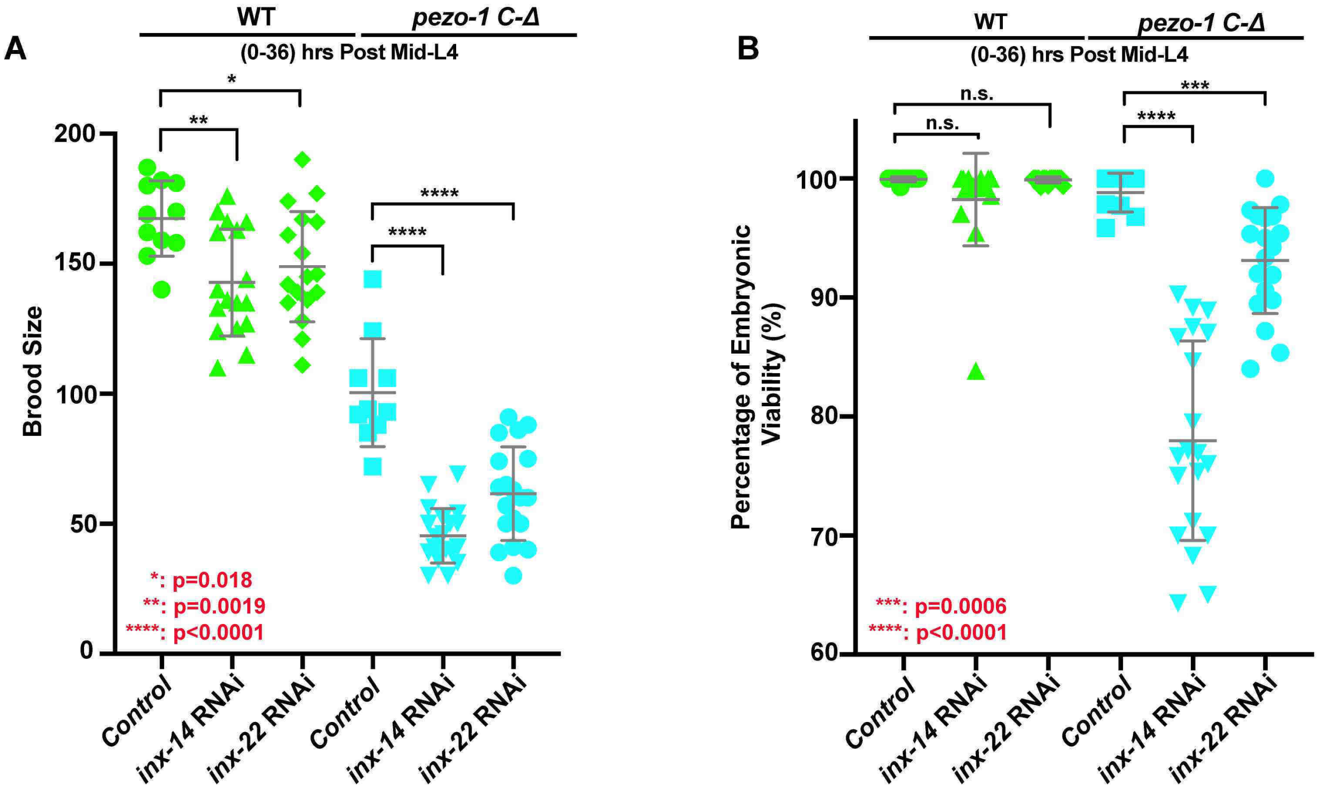

### Supplemental Figure 6

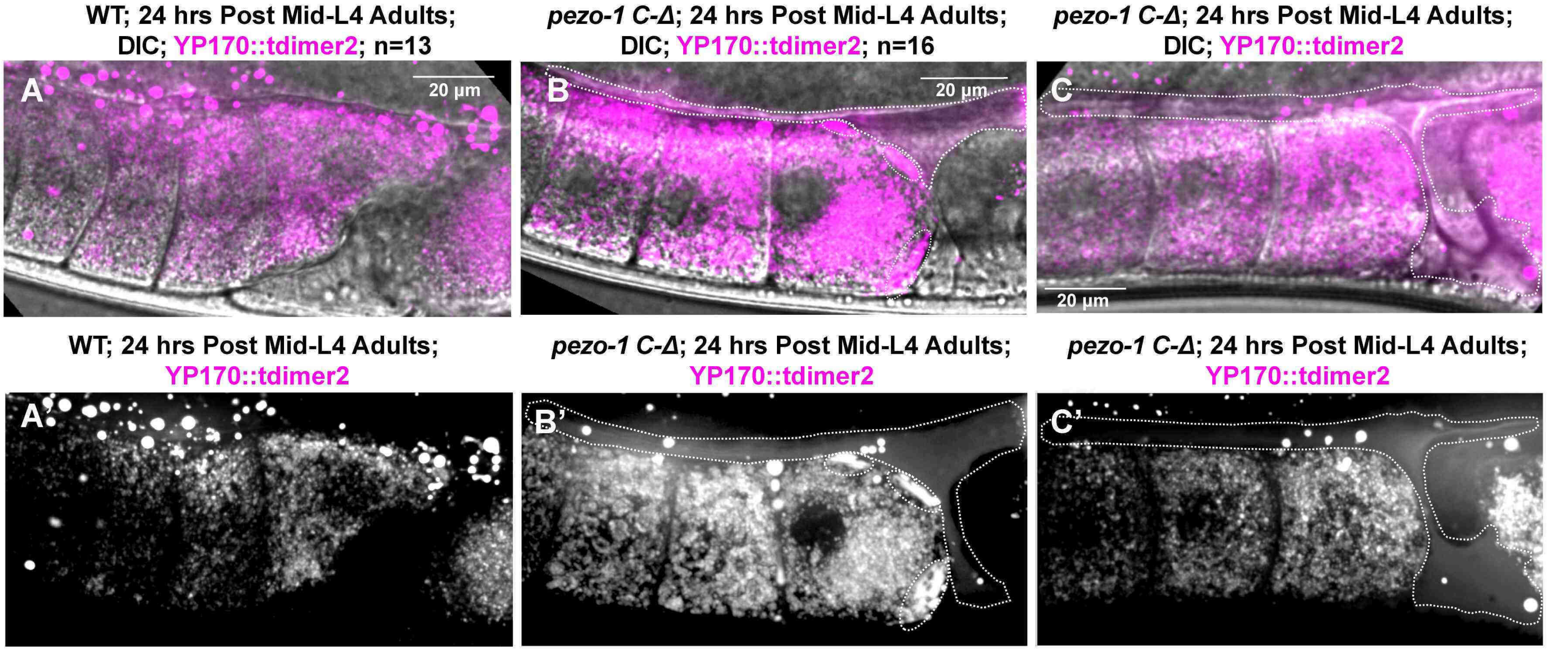
